# The genomic complexity of invasion: cryptic lineages, founder effects, and polygenic sex determination in armored catfish

**DOI:** 10.1101/2025.09.06.674633

**Authors:** Wynne V. Radcliffe, Daemin Kim, Kiedon Bryant, Christopher L. Riggins, Thomas Heard, Monica E. McGarrity, Daniel J. Daugherty, Joshuah S. Perkin, Rachel L. Moran

## Abstract

Genetic biocontrol approaches offer promising tools for managing invasive species. However, their success depends on species identity, demographic history, and the genomic architecture of sex determination, which remain poorly resolved for many non-model taxa. We integrated high-quality genome assembly, whole-genome resequencing, and population genomics to evaluate biocontrol feasibility in invasive suckermouth armored catfish (Loricariidae) in Texas. Mitochondrial phylogenies and genome-wide SNPs revealed cryptic diversity, with distinct lineages from at least two genera (*Hypostomus* and *Pterygoplichthys*) exhibiting strong drainage-level genetic differentiation. Effective population size estimates were exceptionally low, consistent with founder effects and moderate inbreeding. Genome-wide association analyses detected no large-effect loci for sex in either genus, despite high statistical power in *Hypostomus*, suggesting that sex determination is polygenic and/or environmentally influenced. These results identify key biological constraints for sex-ratio-based biocontrol and underscore the importance of genomic assessments prior to management interventions. More broadly, our study highlights how integrating genomic resources, demographic inference, and evolutionary context can guide invasive species control strategies.

## INTRODUCTION

Genetic biocontrol strategies are emerging as powerful tools for managing invasive species, particularly in aquatic ecosystems where traditional methods such as physical removal or chemical treatment can be impractical or ecologically harmful. One promising approach for controlling invasive species involves manipulating sex ratios to promote demographic collapse. In fish species with male heterogametic (XY) sex determination systems, exposure of juvenile males to feminizing hormones can produce phenotypic females that are genetically male (XY “neofemales”). Crosses between neofemales and wildtype XY males are expected to produce 25% YY offspring (“supermales”), which will in turn sire 100% male clutches. This approach has been successful in introduced populations of brook trout (Schill et al. 2016; Schill et al. 2017; Kennedy et al. 2018), Nile tilapia (Gutierrez and Teem 2006), red shiner (Teal et al. 2024), and channel catfish (Wen et al. 2022), with theoretical models and field trials suggesting that repeated introduction of YY supermales can progressively skew the population-level sex ratio towards a male bias and ultimately lead to local extinction.

Despite the potential to be a highly impactful biocontrol approach, the broader applicability of sex-ratio distortion techniques to invasive taxa remains understudied. A major challenge is a lack of key genomic and demographic information in many non-model invasive species. Unresolved taxonomy, unknown population structure, and poorly characterized sex determination systems can all present critical barriers to deploying gene-based control strategies. These knowledge gaps hinder the evaluation and implementation of genetic biocontrol tools in systems where such approaches might otherwise be highly effective. Recent advances in genomic sequencing and analytical tools now make it possible to generate high-resolution population genomic data for non-model species, enabling new insights into invasion dynamics and biocontrol feasibility and providing novel solutions to major challenges in conservation biology.

Suckermouth armored catfishes (family Loricariidae) provide an ideal example for exploring the genomic prerequisites of genetic biocontrol. Introduced through the aquarium trade, these bottom-dwelling catfishes have established dense invasive populations in many rivers, springs, and reservoirs of warm-water climates, particularly Texas and Florida (Jelks et al. 2009). Once introduced, they compete with native species and alter the benthic habitat with their burrowing behaviors. The Texas invasion was first documented in the San Antonio River in the 1960s (Barron 1964), and these catfishes now occur widely, often in ecologically sensitive habitats. These catfishes compete with native species (Scott et al. 2012), alter food web structure (Pound et al. 2011), and cause physical damage to riverbanks through burrowing (Hoover 2004), with significant ecological and economic consequences. In response, local eradication efforts, including spearfishing tournaments and contracted removals, have been initiated in some drainages but have shown limited effectiveness in achieving eradication and their long-term sustainability is undetermined (Blanton et al. 2020; Hay et al. 2022). Moreover, spearfishing targets only visible individuals outside of burrows, and control efforts have so far been uneven across drainages.

Taxonomic uncertainty further complicates management. Morphological (Blanton et al. 2020) and genetic data suggest that species within the genera *Hypostomus* and *Pterygoplichthys* are both present in Texas (Perkin and Bonner 2011), sometimes co-occurring within the same river systems (Arend et al. 2023). This lack of species-level resolution poses a challenge for developing species-specific genetic biocontrol, which requires understanding species boundaries, population connectivity, and sex-determination systems.

This study offers a general framework for using genomic data to evaluate biocontrol feasibility in invasive species with unresolved taxonomy, population structure, and mechanisms of sex determination. Here, we assess the feasibility of genetic biocontrol in invasive suckermouth armored catfish by integrating the first high-quality *de novo* genome assembly for *Hypostomus*, whole-genome resequencing, and population genomic and demographic analyses across multiple drainages in Texas. Our work addresses critical biological prerequisites for gene-based management and provides genomic resources and evolutionary insights that can inform the design and implementation of invasive species control strategies in freshwater ecosystems.

## MATERIALS AND METHODS

### Sample collection and sequencing

Fish were collected via seining and spearfishing from three rivers in central and southern Texas (Texas Parks and Wildlife Department Scientific Permit Number SPR-0218-068). We sampled individuals from the San Marcos River (n = 12 males, n = 11 females) as a part of an annual volunteer spearfishing tournament, San Felipe Creek (n = 12 males, n = 12 females) using seines, and Comal River, from the old channel section only, (n = 11 males, n = 14 females) as a part of contracted spearfishing expeditions coordinated by Atlas Environmental and the Ecological Research Group.

Sex was confirmed via gross dissection of gonads in the field. We dissected white muscle tissue from the lateral side of the body between dorsal and ventral fins for DNA isolation. Whole-body specimens and gonads were preserved in formalin. We extracted DNA from the muscle tissue using the DNeasy Blood and Tissue Kit (Qiagen, Puregene protocol). DNA from one male *Hypostomus* individual from San Marcos River was used for whole genome sequencing (see below) and the remaining 71 samples were sequenced across three lanes of a NovaSeq X Plus 25B flowcell at Texas A&M AgriLife Genomics and Bioinformatics Service.

### Reference genome assembly and annotation

We used DNA from one San Marcos River *Hypostomus* male for reference genome sequencing. A library was prepared using a PacBio HiFi prep kit and was run on one SMRTcell run in HiFi mode on a PacBio Revio machine at the University of Washington Genomics Center. Resulting long-read sequences were assembled using hifiasm (v0.15.1) and wtdbg2 (v2.5) to generate two assemblies from the raw fastq sequencing file (Cheng et al. 2021; Cheng et al. 2022; Cheng et al. 2024; Ruan and Li 2020). We compared the different assembly methods using the benchmarking universal single-copy orthologs (BUSCO) score for genome assembly completeness. In addition, we ran QUAST (v5.2.0) to evaluate the quality of our genome assemblies to assess contiguity, completeness, and accuracy (Gurevich et al. 2013).

We next set out to annotate the reference genome. Prior to annotation, we ran RepeatModeler (v2.0.5) to detect and define repetitive regions in the genome (Flynn et al. 2020). We then ran RepeatMasker (v4.1.4) to functionally mask any repeat regions in the reference genome by replacing their sequence content with N (Tarailo-Graovac and Chen 2009). To determine coordinates in the assembly for genes, exons, and other coding and non-coding sequences, we ran BRAKER3 with AUGUSTUS gene prediction (Stanke et al. 2008; Stanke et al. 2006). Transcriptomic evidence (Gabriel et al. 2024; Brůna et al. 2024; Kovaka et al. 2019; Pertea and Pertea 2020; Quinlan 2014) was incorporated into the genome annotation pipeline from the raw RNA-seq reads for *Pterygoplichthys pardalis* (accession SRR5997830). Reads were downloaded using the fasterq-dump utility from SRA Toolkit (v3.1.1). We also incorporated protein homology evidence to improve gene prediction. Specifically, a curated FASTA file containing protein sequences from a diverse set of fish species was used to guide BRAKER3 and support the identification of conserved gene structures during the annotation process. We then used eggNOG (v5.0) (Huerta-Cepas et al. 2019; Cantalapiedra et al. 2021) to functionally annotate the predicted gene sequences by identifying orthologous genes in the database for Actinopterygii.

### Sequence processing and genotyping

We checked coverage and quality of the raw Illumina whole genome re-sequencing reads for the 71 individuals using FastQC (v0.11.8) (Andrews 2010). We trimmed adapter sequences with barcodes specified for each individual using Cutadapt (v4.2) and Trimmomatic (v0.39) (Martin 2011; Bolger et al. 2014). For downstream analyses, we aligned the processed reads to both the published *Pterygoplichthys pardalis* reference genome (Xia et al. 2022) and our *de novo Hypostomus* sp. genome using bwa (v0.7.17). We used Picard (v2.18.27) to remove duplicates and add read group information. We conducted genotype calling on mapped reads using the Genome Analysis Tool Kit (GATK) following the GATK Best Practices guidelines (McKenna et al. 2010; Van der Auwera et al. 2013; Gabriel and DePristo 2013). First, we used the HaplotypeCaller tool in GATK (v4.5) to generate per-individual gvcfs from the mapped bam files. Then, we used the GenotypeGVCFs tool in GATK (v4.5) to produce vcf files for each scaffold. We applied hard filters using the SelectVariants and VariantFiltration tools in GATK (v4.5). Finally, we used the MergeVcfs tool in GATK (v4.5) to re-combine all subset vcf files for each scaffold. We additionally used BCFtools (v1.19) to only retain biallelic SNPs and remove variants with a minor allele frequency <1% (Danecek et al. 2021).

### Species identification

Previous evidence suggests that the predominant suckermouth armored catfish populations present in the San Marcos River and San Felipe Creek belong to *Hypostomus*, while the predominant species in the Comal River belongs to *Pterygoplichthys* (Gates et al. 2018). However, the species-level identification of suckermouth armored catfish found in Texas rivers remains under debate (see Hay et al. 2022). To help clarify species identity in our whole genome sequencing dataset, we generated consensus mitochondrial gene sequences from variant call format (vcf) files for all samples (Jardim de Quieroz et al. 2020). Mitochondrial genes are particularly useful for species identification due to their high copy number, lack of recombination, and relatively fast mutation rate, which provides sufficient variation to distinguish closely related species. We obtained mitochondrial gene sequences from the densely sampled mitogenomes of Loricariidae and other closely related siluriform species that are publicly available on the National Center for Biotechnology Information (NCBI).

First, we aligned the processed reads to a published *Hypostomus plecostomus* mitochondrial genome (for San Marcos and San Felipe samples) or a *Pterygoplichthys pardalis* mitochondrial genome (for Comal samples), and processed them through the GATK pipeline. This resulted in two vcf files, one for each genus-specific mitochondrial alignment. We then used vcf2fasta and mitochondrial genome annotation files obtained from MitoFish (Zhu et al. 2023) to extract FASTA sequences from the combined vcf for each annotated mitochondrial gene. We extracted mitochondrial gene sequences from each set of samples and used alignments for the cytochrome c oxidase I gene (COI) for species-level identification. COI is known as the standard “barcode” gene for animals and is widely used for this purpose (Andújar et al. 2018). To include data from our reference genome San Marcos River individual, we used MitoHiFi (v3.2) to reconstruct the mitochondrial genome from raw PacBio HiFi sequencing reads. MitoHiFi is a specialized tool that assembles complete, circular mitochondrial genomes by identifying mitochondrial reads from whole-genome sequencing data and assembling them with high accuracy (Uliano-Silva et al. 2023; Allio et al. 2020; Laslett and Canbäck 2008).

We next obtained COI sequence data for other catfishes from NCBI. Following the phylogenetic hypothesis of Lujan et al. (2015), a total of 752 specimens were used in the analysis: 747 loricariids that include *Hypostomus* (n=641), *Pterygoplichthys* (n=74), and several closely related taxa (n=32). Additionally, five *callichthyids* (Corydoras) were included to root the trees (Supplementary Table 8). Phylogenies were inferred with maximum likelihood using IQ-TREE 2 (Minh et al., 2020) from the COI dataset. The best-fit sequence substitution model (TPM2u+F+R5) was determined using ModelFinder (Kalyaanamoorthy et al. 2017) implemented in IQ-TREE 2 based on the Bayesian information criterion (BIC). We assessed topological support with 1000 ultrafast bootstrap replications (Hoang et al. 2018).

### Population genetics

We conducted population structure analyses in two steps. First, we aligned all samples to the same reference genome of *Hypostomus plecostomus* to identify patterns in and across all samples. We then conducted the same analyses on only “true” *Hypostomus* species, the San Felipe Creek and San Marcos River populations, to elucidate differences in species-level gene flow. To assess genetic variation among populations, we conducted a principal component analysis (PCA) using PLINK v2.00a3.7. For the PCA, we used alignments for the entire dataset to the *Hypostomus* reference genome and applied linkage disequilibrium (LD) pruning to reduce the effects of linked markers and retain only independent SNPs. We used the software ADMIXTURE (v1.3.0) to further assess population structure and gene flow (Alexander et al. 2009). ADMIXTURE is a maximum likelihood estimation tool for inferring individual ancestries from multilocus SNP genotype datasets. Analyses were performed on the PLINK-formatted binary genotype file generated for the PCA analysis (Chang et al. 2015). We ran ADMIXTURE for values of K (the number of predicted ancestral populations) ranging from 1 to 5 (Supplementary Figure 2). Each run was performed with 10-fold cross-validation (CV). The optimal number of clusters (K) was determined by identifying the value with the lowest CV error (Supplementary Table 9), or the point at which the CV error showed the greatest decrease compared to the previous K value, indicating improved model fit with additional clusters.

We next wanted to use our genetic data to obtain an estimate of effective population size (Ne) for each population of suckermouth armored catfish in our dataset, to compare to previous mark recapture-based estimates (Hay et al. 2022). We used currentNe (Santiago et al. 2024) to estimate a contemporary effective population size for the San Marcos River and San Felipe Creek samples aligned to the *Hypostomus* reference genome and the Comal River samples aligned to the *Pterygoplichthys* reference genome. We randomly sampled 1,000,000 SNPs from our dataset as an input file for the estimation to comply with the program’s limitations. Preliminary results indicated that effective population sizes were much smaller than previously estimated census sizes (C. Riggins, personal communication, April 2, 2025), suggesting high levels of inbreeding. This is consistent with the expectation that these introduced populations were founded by a small number of individuals from the aquarium trade. Because inbreeding reduces genetic variation, estimates of effective population size based on genetic diversity metrics are likely to be downwardly biased.

Runs of homozygosity (ROH) can also be used as a metric for estimating inbreeding and effective population size, as they reflect regions of the genome where individuals have inherited identical segments from both parents. These segments exceed levels of homozygosity that would be expected under Hardy-Weinberg equilibrium. Populations with longer ROH typically have a history of more prolonged or recent inbreeding, while shorter ROH may indicate older or less intense inbreeding events. We used RZooRoH (Druet and Gautier 2017; Bertrand et al. 2019) to estimate inbreeding and detect homozygous-by-descent (HBD) segments in our study populations. RZooRoH calculates the lengths of HBD segments for each individual in a population and uses a model to estimate the age of each segment based on its length. The model we used separates HBD segments into age-related classes and estimates the contribution of these classes to the current levels of homozygosity seen in a population. A larger contribution by a class to current homozygosity indicates a reduced effective population size at the class’ corresponding period, possibly associated with bottleneck events or founder effects. More segments of homozygosity (i.e., a higher contribution value) are expected if populations are small and experience genetic drift and inbreeding. Conversely, low contribution by a class indicates a higher effective population size at that time. Previous estimates of effective population size for the San Marcos River *Hypostomus* population have come from N-mixture modelling of video camera capture data (Byckovski et al. 2023). Therefore, we only ran RZooRoH on the individuals we had genomic data for from the San Marcos population to compare to the previous video-based estimates.

To further assess the level of inbreeding within each population, we calculated inbreeding coefficients (F_IS_) for each population. We used VCFtools (v0.1.16) to calculate the F_IS_ statistic, which is based on observed and expected homozygosity in a population. The command vcftools --het was applied to the filtered vcf files for each population. This command outputs a per-individual inbreeding coefficient following the formula 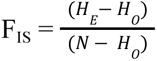 in which H is the number of homozygous genotypes expected under Hardy-Weinberg Equilibrium, H_O_ is the number of homozygous genotypes observed in our data, and N is the total number of genotyped sites analyzed in our data. This coefficient reflects deviation from Hardy-Weinberg proportions within individuals and serves as an estimate of the inbreeding coefficient of an individual relative to its subpopulation. We then averaged all F_IS_ values across the population to estimate the mean inbreeding coefficient for pooled males and females.

### Investigating genetic sex determination

We performed a genome-wide association study (GWAS) to identify genomic regions associated with sex using linear mixed models in GEMMA (v0.98.5) (Zhou and Stephens 2012). These analyses were conducted on samples aligned to their most closely related reference genome in order to minimize variation attributed to reference bias. We first estimated a centered relatedness matrix using the -gk 2 option from PLINK-formatted genotype data to account for population structure and relatedness among individuals. The resulting matrix was used as input for a linear mixed model association test, using the -lmm 4 option, which computes Wald, likelihood ratio, and score test statistics for each SNP. We analyzed a total of 5,073,283 SNPs in the pooled set of San Marcos River and San Felipe Creek *Hypostomus* populations (n = 23 males and n = 23 females) and 480,206 SNPs from the Comal River *Pterygoplichthys* population (n = 11 males, n = 14 females).

We estimated statistical power for a GWAS of sex using a standard logistic-test approximation at a two-sided genome-wide threshold of α = 5×10^-8^. For a fully sex-linked, non-recombining locus in an XX/XY or ZZ/ZW system, when sexes are sampled in equal numbers one sex is homozygous and the other heterozygous, we expect a pooled MAF ≈ 0.25. We therefore calculated the odds ratio (OR) required to reach 80% power at MAF = 0.25 for each dataset.

## RESULTS

### Genome assembly and annotation

We used two approaches to generate genome assemblies from long-read sequencing data for a single male specimen collected from the San Marcos River. The wtdbg2 assembly consisted of 5,620 contigs, with a total genome size of 1.371 Gb, an N50 of 5.98 Mb, and a GC content of 40.45%. In comparison, the hifiasm assembly consisted of 2,267 contigs, with a total genome size of 2.077 Gb, an N50 of 7.6 Mb, and a GC content of 40.63%. The higher N50 and lower contig count in the hifiasm assembly suggest a more contiguous and complete genome reconstruction relative to the wtdbg2 assembly. Furthermore, a previous estimate based on flow cytometry also indicated an expected genome size of approximately 2 Gb for *Hypostomus* (https://www.genomesize.com), providing further support that the hifiasm assembly is more complete relative to the wtdbg2 assembly.

To further evaluate assembly completeness, we performed a BUSCO (Benchmarking Universal Single-Copy Orthologs) analysis. The wtdbg2 assembly had a BUSCO completeness score of 96.8% (C: 96.8% [S: 96.0%, D: 0.8%], F: 0.6%, M: 2.6%), while the hifiasm assembly scored 98.1% (C: 98.1% [S: 63.0%, D: 35.1%], F: 0.7%, M: 1.2%). Based on both structural metrics and completeness, the hifiasm assembly was selected as the reference genome for all downstream analyses.

RepeatModeler found and classified 2,308 repeat families from the hifiasm genome assembly, using 61.24% of the assembly to uncover the repeats. Of the known families, DNA/TcMar-Tc1, a class and superfamily of interspersed repeats and DNA transposons, were the most represented with 187 repeat segments identified through RepeatModeler (Supplementary Table 7). RepeatMasker masked 62.23% of the base pairs in the hifiasm genome assembly by replacing them with “N” characters. According to the output report, 60.01% of the genome consisted of interspersed repeats, all of which were unclassified, while an additional 2.23% comprised simple repeats. In both reference assemblies, the most represented repeat families were interspersed DNA elements.

### Whole genome re-sequencing

We assessed sequence quality at each stage of the data processing pipeline using FastQC and quantified read retention and coverage throughout the alignment workflow. Initial raw fastq files showed high base quality scores across all reads. After preprocessing (e.g., Trimmomatic and CutAdapt), FastQC confirmed that read quality remained high, with no base pairs being lost in between processing steps (Supplementary Tables 1-6). Once we aligned each populations’ reads to their respective reference genome, we monitored coverage and read mapping statistics of the alignment files at each processing stage. Mean coverage remained consistently high across samples (Supplementary Tables 1-6), with minimal loss during sorting, deduplication, or indexing. Final mean ± SE coverage of each population after mapping was 14.40 ± 4.62 for San Marcos River, 17.22 ± 7.96 for the Comal River, and 12.23 ± 4.80 for the San Felipe Creek samples.

### Species identification

Results of the phylogenetic analysis using 752 COI sequences of loricariid and callichthyd specimens identified the San Marcos River reference and both San Felipe Creek individuals as the sister lineage to *Hypostomus robinii*, a species endemic to Trinidad (Figure 1). The reference sequence used for *H. robinii* was derived from a specimen collected in Trinidad (Jardim de Queiroz et al. 2020; see Supplementary Table 8), suggesting our sequences are accurately assigned at the species level. The Comal River individual was nested within a *Pterygoplichthys* clade, alongside several *Pterygoplichthys pardalis* sequences and putative *Hypostomus plecostomus* individuals (Figure 2).The putative *H. plecostomus* specimens that were placed in a *Pterygoplichthys* clade have an aquarium trade origin (see Supplementary Table 8), which suggests that they were potentially misidentified *P. pardalis* (as “*H. plecostomus*”), as species misidentification is not uncommon in aquarium trades (Collins et al. 2012).

**Figure 1:**
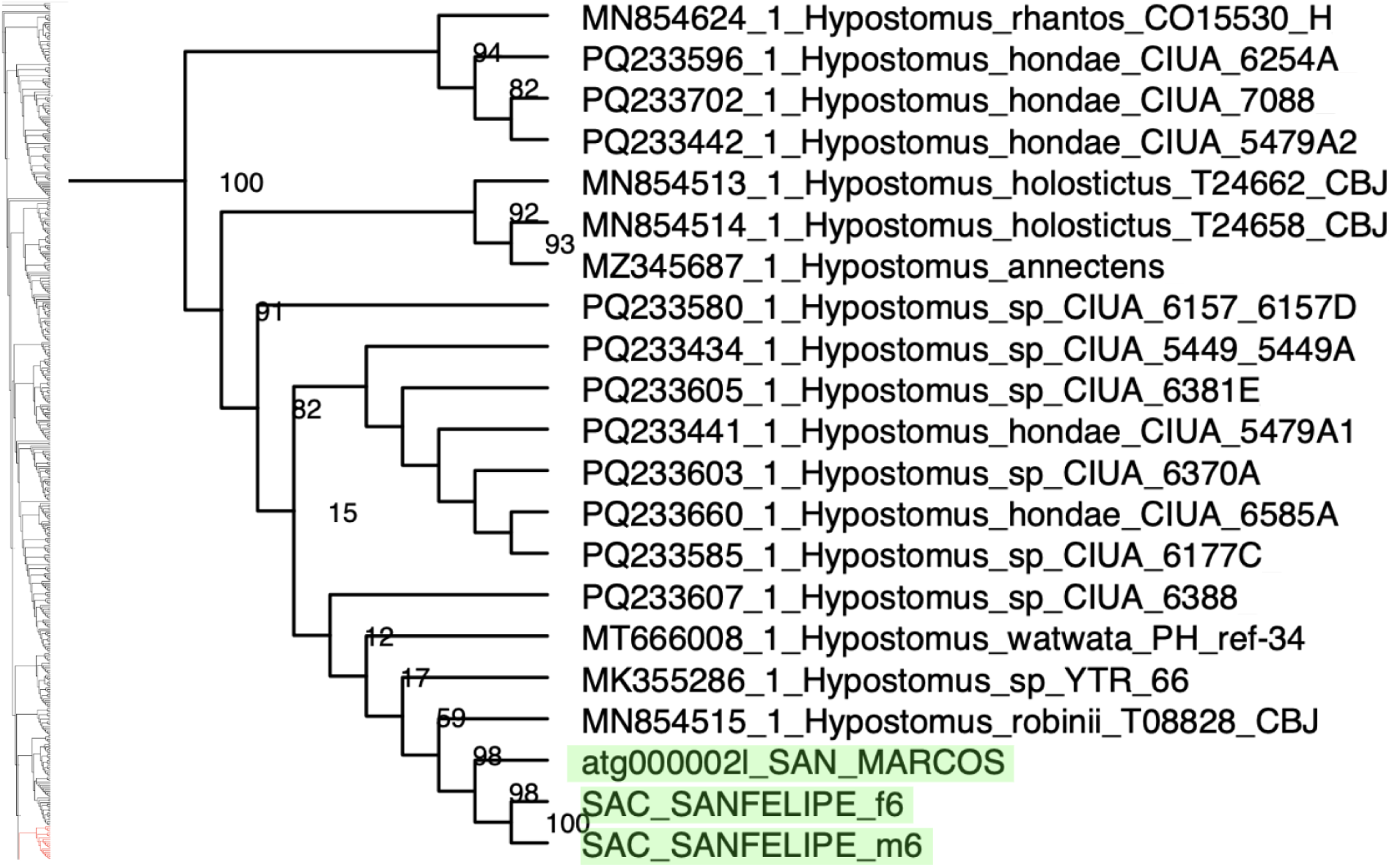
Results of IQ-TREE2 analysis showing the phylogenetic hypotheses of the clade that includes San Marcos River and San Felipe Creek populations of suckermouth armored catfish (highlighted in green). Numbers on the nodes indicate bootstrap support values. To see the position of this clade within the complete COI gene tree, see Supplementary Figure 1.

**Figure 2:**
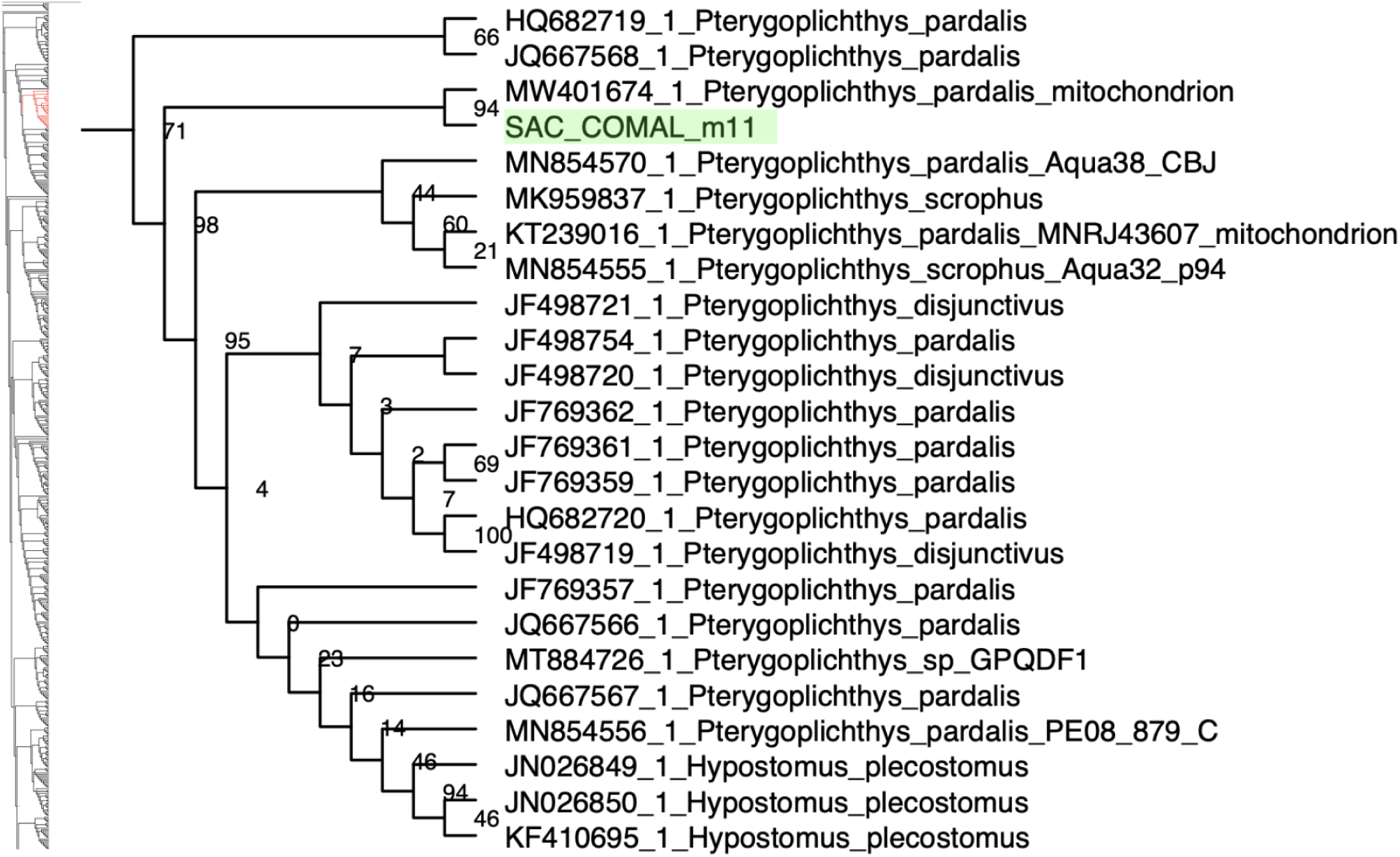
Results of IQ-TREE2 analysis showing the phylogenetic hypotheses of the clade that includes the Comal River population of suckermouth armored catfish (highlighted in green). Numbers on the nodes indicate bootstrap support values. To see the position of this clade within the complete COI gene tree, see Supplementary Figure 1.

### Population structure

We performed principal component analysis (PCA) on SNP genotype data to assess genetic structuring among populations from Comal, San Felipe, and San Marcos rivers. When we conducted the PCA on all populations aligned to the *Hypostomus* reference genome, we found evidence for strong genetic differentiation by population. The first principal component (PC1) explains 44.05% of the total genetic variation, clearly separating individuals by population (Figure 3a). We observed the same strong population structure when we restricted our PCA to only the “true” *Hypostomus* populations – those that the mitochondrial phylogeny identified as such. The first principal component (PC1) explains 18.47% of the total genetic variation and clearly separates individuals by population, indicating strong population structure (Figure 3b). PC2 explains 9.35% of the variation, but does not appear to correlate with known factors like sex or geography. To determine the significance of these associations, we then performed a PERMANOVA test based on the main principal components, PC1 and PC2, for each set of analyses. For all samples aligned to the *Hypostomus* reference genome, population explains 96% of the variation in SNP data (R² = 0.961), with significant statistical support (F = 760, p = 0.001). This indicates that genetic clustering is almost completely determined by population, with very little overlap between populations of suckermouth armored catfish across genera. Individuals from each river form distinct clusters with no overlap, suggesting limited gene flow and strong genetic differentiation among populations. Similar results were observed when restricting to the “true” *Hypostomus* groups in the San Marcos River and San Felipe Creek. In these samples, population explains 48% of the variation in SNP data (R² = 0.477, F = 40.1, p = 0.001).

**Figure 3:**
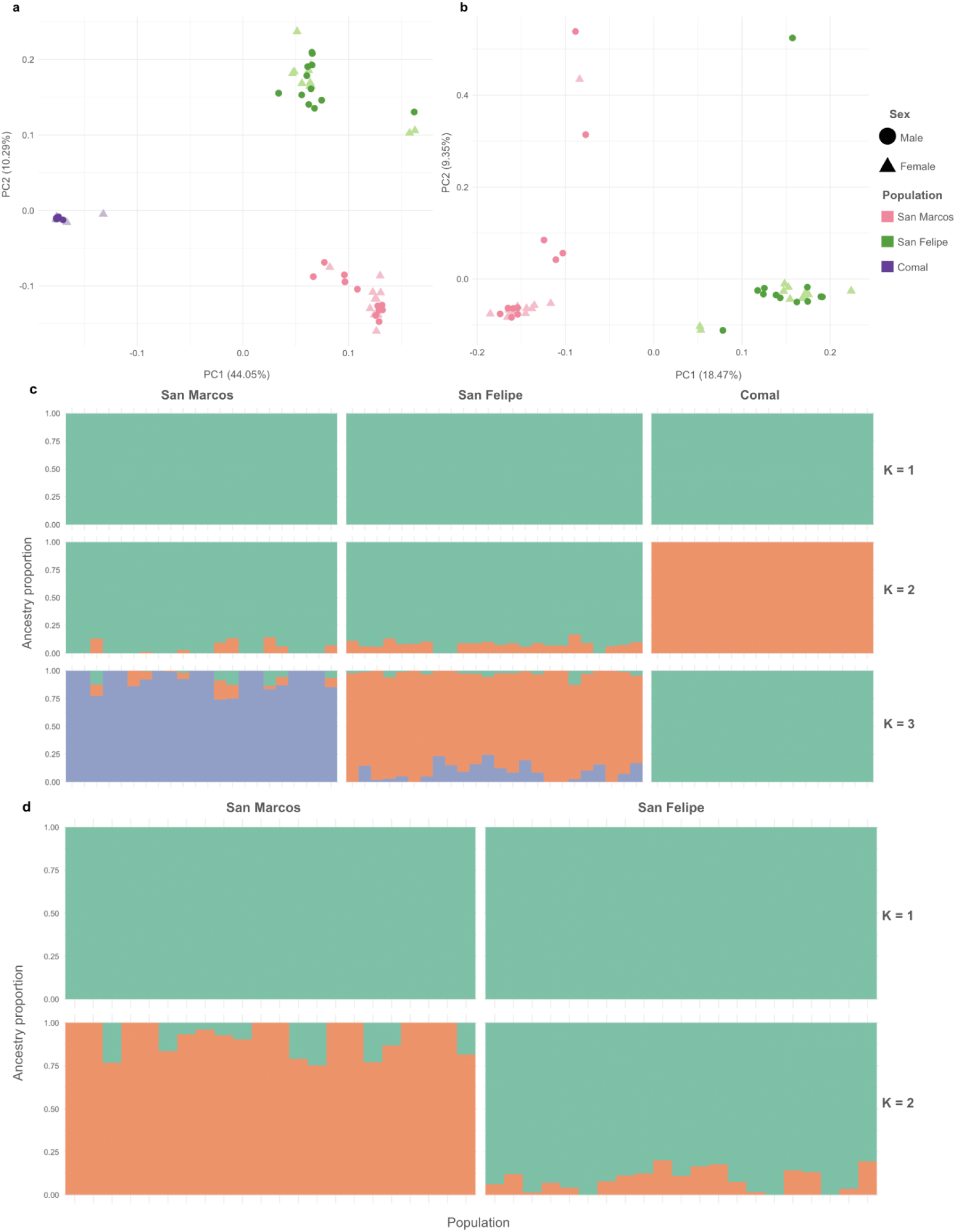
(a) Principal component analysis (PCA) plot of all samples aligned to the *Hypostomus* reference genome. Shapes indicate sex, with triangles representing females and circles representing males. (b) PCA plot showing distinct genetic variation among the two *Hypostomus* populations. (c) Genetic population structure of suckermouth armored catfish using ADMIXTURE with all three Texas rivers shows historical, but not contemporary, gene flow. The optimal number of clusters (K=3) indicates slight shared ancestry among all three populations. (d) ADMIXTURE analysis only including the two true *Hypostomus* populations showed similar patterns, with the most supported number of clusters (K=2) showing some degree of shared ancestry between the San Felipe Creek and San Marcos River.

Results of our ADMIXTURE analysis of genetic population structure identified K = 3 as the most accurate clustering value for the all-population dataset (Supplementary Table 9). Samples from the Comal River population showed no evidence of genetic material from the San Felipe Creek or San Marcos River samples (Figure 3c), consistent with their classification as a different genus. We next limited our population structure analysis to the San Marcos River and San Felipe Creek populations because of their close phylogenetic relationships. Results indicate limited shared genetic ancestry between the two populations, though they remain largely distinct clusters (Figure 3d). Based on the cross-validation error rate, K = 2 was the most accurate clustering value for this two-population dataset (Supplementary Table 9).

### Inbreeding and genetic diversity

Our effective population size analysis with currentNe yielded surprisingly low values. For the San Marcos River and San Felipe Creek populations, currentNe mean ± SE estimates were relatively close, 63.60 (±10.26) and 63.20 (± 10.22), respectively, but the Comal River population estimate was slightly higher at 81.20 (± 17.52). After averaging individual inbreeding coefficients (F_IS_), we found mean ± SE values of 0.031(± 0.046) for the San Marcos River population, 0.094 (± 0.059) for the San Felipe Creek population, and 0.032 (± 0.104) for the Comal River population. F_IS_ values range from -1 to 1: values above 0 indicate more homozygosity than expected under Hardy-Weinberg equilibrium, suggesting inbreeding, while values below 0 suggest excess heterozygosity in the population. We next estimated homozygous-by-descent (HBD) segments for three age-related classes (K = 3). For K = 3, the contribution of classes 1 and 3 was zero, while class 2 had a 100% contribution to the segments. All segments in the San Marcos River population best match a time frame consistent with a historical inbreeding event.

### Sex determination mechanism

Our analysis did not identify any SNPs across either the *Hypostomus* or *Pterygoplichthys* samples that were significantly sex-associated (Figure 4). Similar patterns were found when we grouped San Marcos River and San Felipe Creek populations (both *Hypostomus* species) or analyzed each population independently. No significant peaks were found across linkage groups for either species. Given our sample sizes and an expected MAF = 0.25 and α = 5×10^-8^, achieving 80% power required OR = 20.7 in *Hypostomus* and OR = 108.9 in *Pterygoplichthys* (Figure 4c). Here, OR quantifies how strongly genotype predicts phenotypic sex; larger OR indicates a more deterministic genotype–sex relationship. These thresholds indicate our study is powered to detect only strong, large-effect loci associated with sex, such as those expected for fully sex-linked markers.

**Figure 4:**
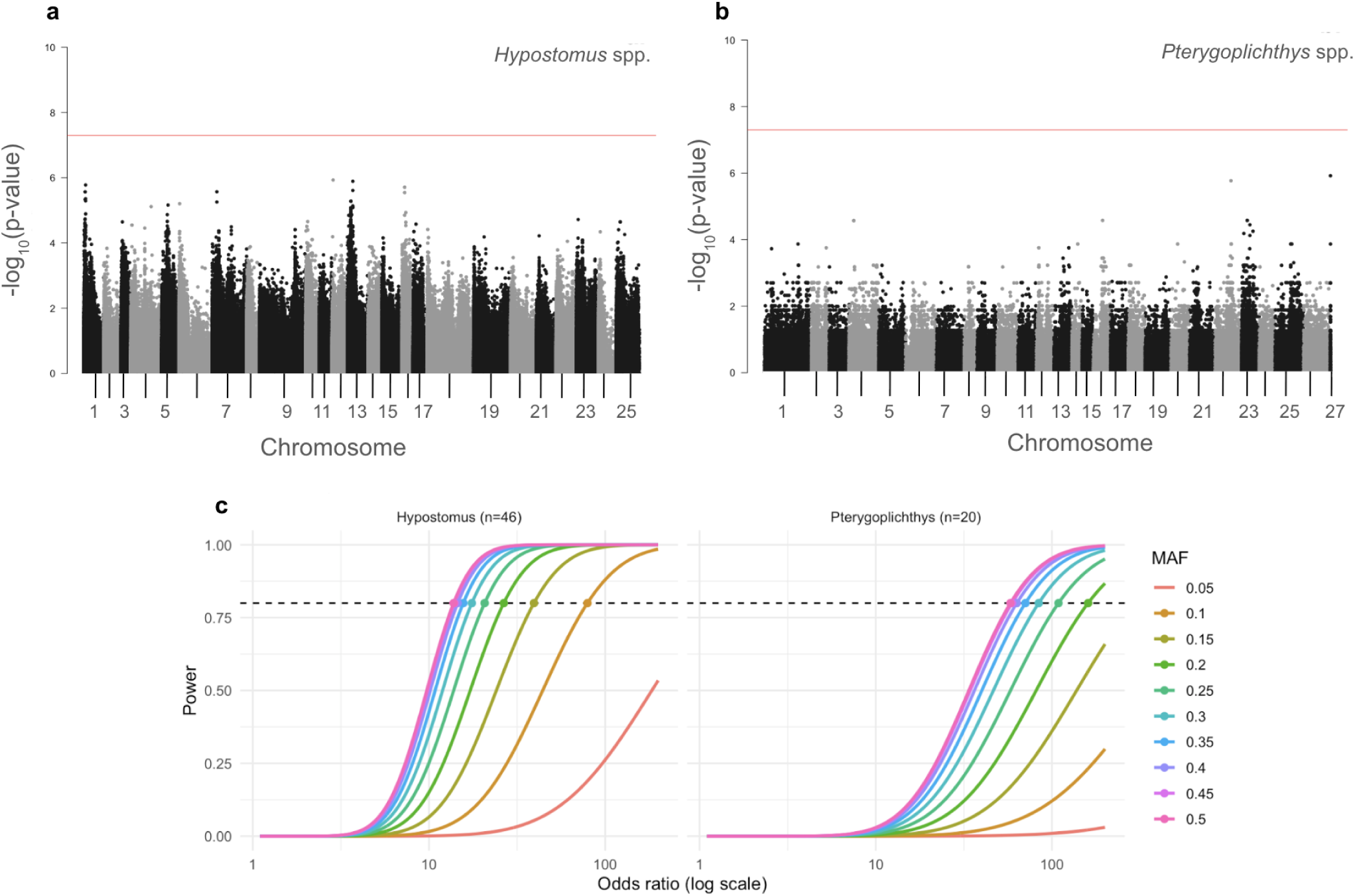
Genome-wide association study using sex as the phenotype showed no single region of the genome where sex is more differentiated than another. Samples from the San Marcos River and San Felipe Creek (n = 23 males and n = 23 females) aligned to the *Hypostomus* reference genome (a) and samples from the Comal River (n = 11 males and n = 14 females) aligned to the *Pterygoplichthys* genome (b). The genome-wide significance threshold (p = 5e-08) is shown as a horizontal red line. A lack of significantly sex-associated SNPs (above the threshold) suggests the sex-determining system of suckermouth armored catfish is likely to be environmental or polygenic. (c) Predicted power to detect sex-associated loci across effect sizes in *Hypostomus* and *Pterygoplichthys* GWAS analyses. Curves shown for MAF = 0.05 to 0.50 at and α = 5×10^-8^.

## DISCUSSION

Our study demonstrates how integrating genomic resources, demographic inference, and evolutionary context can clarify the feasibility of genetic biocontrol in non-model invasive species. By combining a high-quality reference genome, phylogenetic analysis, and population-level resequencing, we found that invasive suckermouth armored catfish in Texas comprise multiple cryptic lineages with strong genetic differentiation among drainages, low effective population sizes, and a likely polygenic or environmentally influenced basis of sex determination. These findings highlight key biological constraints that could preclude the use of sex ratio-based biocontrol, yet also identify demographic features that may increase vulnerability to targeted management.

The high-contiguity *Hypostomus* genome we generated provides an essential platform for future genetic and comparative studies in Loricariids. The large genome size and high repeat content of our *Hypostomus* reference genome was similar to that of the published *Pterygoplichthys pardalis* chromosome-level assembly (Xia et al. 2022). This repeat-rich structure parallels patterns seen in other teleosts with complex genome histories (Braasch and Postlethwait 2012; Inoue et al. 2015; Blanco et al. 2014; Glasauer and Neuhauss 2014), and provides an essential resource for investigating genome evolution in Loricariids, including the role of transposable elements in shaping structural variation and adaptive potential. Repetitive elements, particularly transposable elements, are well known to drive genome restructuring and regulatory rewiring in fishes (Braasch and Postlethwait 2012; Inoue et al. 2015). For example, bursts of DNA transposons and LINE elements have been linked to chromosomal rearrangements that promote local adaptation in cichlids and killifishes (García et al. 2022, Almeida et al. 2025). Invasive species often experience strong bottlenecks, yet their success is thought to depend on genomic mechanisms that generate novel variation, including mobilization of transposable elements under stress (Stapley et al. 2015). Mapping the distribution and activity of repeats in *Hypostomus* will therefore be critical for testing whether repeat-associated structural variants contribute to invasion success in armored catfish. From an applied perspective, high repeat content also poses challenges for designing CRISPR-based or gene-drive biocontrol strategies, as repetitive sequences can lead to widespread off-target effects (Naeem et al. 2020). Understanding the repeat landscape in *Hypostomus* will thus inform both evolutionary models of invasion and the technical feasibility of genome-targeted management interventions.

Phylogenetic analyses identified the San Marcos River and San Felipe Creek populations as closely related to *Hypostomus robinii*, contradicting earlier taxonomic assignments (Pound et al. 2011). The Comal River population clustered within *Pterygoplichthys*, corroborating previous claims that the species in that river was *Pterygoplichthys disjunctivus* (Hay et al. 2022). This taxonomic clarification is critical because mismatched management strategies across lineages could undermine control efforts. Our PCA and ADMIXTURE analyses robustly grouped samples from the same population together and showed separation by genus, supporting the results of our mitochondrial gene tree (i.e., San Marcos River and San Felipe Creek *Hypostomus* spp. clustered closer together, Comal River *Pterygoplichthys* spp. more distant). These patterns are consistent with inbreeding in isolated, introduced populations.

Indeed, demographic analyses revealed founder effects, moderate inbreeding, and historically reduced effective population sizes across populations, likely reflecting introduction via the aquarium trade followed by limited dispersal. Levels of homozygosity suggest an intermediate-level event such as a population bottleneck in the last few generations. Previous video surveys conducted at the Meadows Center in San Marcos, Texas, estimated a census population (Nc) size ranging from 5,000 to 12,000 depending on the season (Perkin & Heard 2023), approximately 80 - 200X larger than our estimates based on genome-wide SNP data. We suspected that our apparent underestimate could be influenced by inbreeding depression, which reduces the pool of available genetic variation used to infer Ne. The San Marcos River population showed consistent inbreeding signals from the intermediate heterozygosity-by-descent class, which might reflect a historic but not ancient collapse in effective population size. This dominance is consistent with a classic founder effect: a population that was initially founded by a small number of individuals, likely introduced through the pet trade. Additionally, periodic geographic barriers, such as river freezes and droughts, are known to occur in this system and may temporarily restrict gene flow, particularly among subpopulations within the San Marcos River or San Felipe Creek system. If these events happen cyclically, they could act as recurring bottlenecks, reinforcing the signal of historical inbreeding and contributing to the observed patterns of homozygosity over time.

Similar demographic signatures have been observed in other successful aquatic invaders, such as lionfish in the western Atlantic (Bernardi et al. 2024), where reduced genetic diversity did not prevent establishment but shaped invasion trajectories. In armored catfish, low Ne may likewise constrain adaptive potential while increasing susceptibility to targeted control measures (Chiesa et al. 2019). Strong differentiation among drainages and lineage-specific demographic histories further emphasize that management strategies must be tailored to local population dynamics, as has been shown in other aquatic invaders with restricted gene flow (Barthel et al. 2010; Sorensen and Hoy 2007). The San Marcos River, for instance, has had significant suppression efforts since at least 2013, with over 16,000 fish removed (Blanton et al. 2020). Our results suggest that low genetic diversity and evidence of repeated bottlenecks in this population could make it more vulnerable to removal or environmental stressors, whereas deeply divergent lineages may differ in traits such as fecundity, growth, or habitat tolerance that influence management outcomes. Recognizing this genomic and taxonomic complexity is therefore essential for designing effective interventions, whether through physical removal, habitat modification, or the evaluation of genetic biocontrol strategies.

Our genome-wide association analyses detected no large-effect sex-linked loci in either genus, despite high power to detect such loci in *Hypostomus* and moderate power in *Pterygoplichthys*. This points to a polygenic or environmentally influenced sex-determination system, consistent with the evolutionary lability of sex determination in teleosts but contrasting with other armored catfish that exhibit clear chromosomal sex determination (Blanco et al. 2014; Balini et al. 2024). While our study design had sufficient power to identify major-effect loci, smaller-effect variants or environmentally mediated mechanisms may have gone undetected. Broader sampling and controlled breeding experiments will be essential to disentangle genetic from environmental contributions to sex determination, and to evaluate whether intermediate-effect loci could still be leveraged for biocontrol. These findings underscore that lineage-specific sex-determination systems may limit the feasibility of sex ratio–distortion strategies in invasive armored catfish, highlighting the need for diversified management approaches.

Overall, our results underscore that invasive armored catfish in Texas represent a mosaic of cryptic lineages with distinct demographic histories and complex sex-determination systems, that may require nuanced, lineage-specific management approaches. More broadly, our work illustrates the value of genomic assessments before implementing genetic biocontrol, particularly in taxa with uncertain taxonomy, limited connectivity, and poorly understood mechanisms of sex determination. Beyond armored catfish, this approach can inform feasibility assessments for other invasive species with uncertain taxonomy, poorly resolved population structure, and uncharacterized sex determination systems.

Strengthening regulations on aquarium trade, combined with genomically informed management, will be key to reducing the ecological impacts of non-native freshwater fishes.

## Author Contributions

DJD and MM assisted with experimental design and project planning. KB, JSP, CLR, and TH collected samples from the Comal River, San Felipe Creek, and San Marcos River. KB performed all DNA extractions for sequencing. RLM was responsible for the reference genome assembly. WVR conducted population genomics and sex association analyses and was responsible for all data processing. DK performed phylogenetic analyses on mitochondrial data for species identification. All authors contributed to writing the manuscript.

## Acknowledgements

We thank the TxGen Genomics Core for their guidance and performing the 10X library preparations and Illumina sequencing. The treatment of animals used in this study was in compliance with Texas A&M University’s Institutional Animal Care and Use Committee (IACUC) under AUP #8902. The Texas A&M University HPRC provided resources that contributed to the research results reported within this paper. We thank Emily Lorkovic and Collin Garoutte for their help in collecting individuals for this project as well as providing informative ecological data. Fish were collected under the Texas Parks and Wildlife Department under Scientific Permit Number SPR-0218-068. The Meadows Center for Water and the Environment at Texas State University as well as Atlas Environmental helped us to procure fish that were speared as a part of routine research or spearfishing control programs.

## Disclosure

Data Accessibility: All raw reads have been deposited to NCBI SRA (PRJNA1198926) and will be released upon publication. Scripts for data processing and analysis are available at https://github.com/evoradcliffe/SAC_TPWD_WGS.

Benefits Generated: Benefits from this research accrue from the sharing of our data and results on public databases as described above.

## Data Availability Statement

All raw reads have been deposited to NCBI SRA (PRJNA1198926) and will be released upon publication. Scripts for data processing and analysis are available at https://github.com/evoradcliffe/SAC_TPWD_WGS.

## Supporting Information

Additional supporting information can be found online in the Supporting Information section.

