## Supplemental Information for "The genomic complexity of invasion: cryptic lineages, founder effects, and polygenic sex determination in armored catfish"

### Table of Contents:


| | |
|---|---|
| **Table 1: San Marcos female sequencing stats** | Page 2 |
| **Table 2: San Marcos male sequencing stats** | Page 3 |
| **Table 3: Comal female sequencing stats** | Page 4 |
| **Table 4: Comal male sequencing stats** | Page 5 |
| **Table 5: San Felipe female sequencing stats** | Page 6 |
| **Table 6: San Felipe male sequencing stats** | Page 7 |
| **Table 7: Repeat families** | Pages 8-10 |
| **Table 8: COI sequence metadata** | Pages 11-36 |
| **Table 9: CV error rates** | Page 37 |
| **Figure 1: Full COI phylogeny** | Page 38 |
| **Figure 2: ADMIXTURE plots for K=1 to K=5** | Page 39 |

*Supplementary Table 1:* Sequencing statistics and quality for n = 11 San Marcos female samples at each stage of the data processing pipeline for raw reads as well as coverage throughout the alignment workflow. Coverage statistics are based on an alignment to *Hypostomus* reference genome.

| | F1 | F2 | F3 | F4 | F5 | F6 | F7 | F8 | F9 | F10 | F11 |
|---|---|---|---|---|---|---|---|---|---|---|---|
| **R1** | | | | | | | | | | | |
| Raw (Lane 1) | 57270881 | 54601736 | 50423678 | 52117557 | 44115779 | 56806880 | 45036518 | 47645233 | 74228760 | 54108225 | 70935253 |
| Raw (Lane 2) | 57417123 | 54442246 | 49893146 | 51290629 | 44653061 | 56513183 | 44685820 | 47382193 | 73530669 | 53234847 | 70666892 |
| Raw (Lane 3) | 57292580 | 54557315 | 50801311 | 51935584 | 43250105 | 56790951 | 45252923 | 47171993 | 74142678 | 53742428 | 71631978 |
| Raw Total | 171980584 | 163601297 | 151118135 | 155343770 | 132018945 | 170111014 | 134975261 | 142199419 | 221902107 | 161085500 | 213234123 |
| Concatenated | 171980584 | 163601297 | 151118135 | 155343770 | 132018945 | 170111014 | 134975261 | 142199419 | 221902107 | 161085500 | 213234123 |
| CutAdapt | 171980584 | 163601297 | 151118135 | 155343770 | 132018945 | 170111014 | 134975261 | 142199419 | 221902107 | 161085500 | 213234123 |
| Trimmomatic Out (R1) | 153746590 | 143240504 | 133215320 | 137987946 | 120713082 | 150949144 | 119205556 | 127774380 | 199338553 | 141798864 | 189678708 |
| **R2** | | | | | | | | | | | |
| Raw (Lane 1) | 57270881 | 54601736 | 50423678 | 52117557 | 44115779 | 56806880 | 45036518 | 47645233 | 74228760 | 54108225 | 70935253 |
| Raw (Lane 2) | 57417123 | 54442246 | 49893146 | 51290629 | 44653061 | 56513183 | 44685820 | 47382193 | 73530669 | 53234847 | 70666892 |
| Raw (Lane 3) | 57292580 | 54557315 | 50801311 | 51935584 | 43250105 | 56790951 | 45252923 | 47171993 | 74142678 | 53742428 | 71631978 |
| Raw Total | 171980584 | 163601297 | 151118135 | 155343770 | 132018945 | 170111014 | 134975261 | 142199419 | 221902107 | 161085500 | 213234123 |
| Concatenated | 171980584 | 163601297 | 151118135 | 155343770 | 132018945 | 170111014 | 134975261 | 142199419 | 221902107 | 161085500 | 213234123 |
| CutAdapt | 171980584 | 163601297 | 151118135 | 155343770 | 132018945 | 170111014 | 134975261 | 142199419 | 221902107 | 161085500 | 213234123 |
| Trimmomatic | 153746590 | 143240504 | 133215320 | 137987946 | 120713082 | 150949144 | 119205556 | 127774380 | 199338553 | 141798864 | 189678708 |
| **Total # Reads and Base Pairs** | | | | | | | | | | | |
| Total reads | 169494559 | 160818662 | 148450428 | 152299520 | 130212374 | 167401005 | 132608474 | 139803052 | 217861776 | 158016197 | 209820673 |
| Total base pairs | 48486172350 | 45608874900 | 42249862200 | 43543119900 | 37638818400 | 47752522350 | 37772104500 | 40136614800 | 62580049350 | 44972259150 | 59924907150 |
| **Coverage** | | | | | | | | | | | |
| Pre-alignment | 24.243086175 | 22.80443745 | 21.1249311 | 21.77155995 | 18.8194092 | 23.876261175 | 18.88605225 | 20.0683074 | 31.290024675 | 22.486129575 | 29.962453575 |
| Post-alignment | 21.9566 | 20.3221 | 19.1833 | 19.7845 | 16.8709 | 21.5172 | 17.1483 | 18.2744 | 28.33 | 20.3378 | 27.043 |
| Rgadd | 21.9566 | 20.3221 | 19.1833 | 19.7845 | 16.8709 | 21.5172 | 17.1483 | 18.2744 | 28.33 | 20.3378 | 27.043 |
| Dupsmarked | 12.0193 | 12.9093 | 11.8424 | 12.6621 | 12.0495 | 13.7355 | 10.2776 | 12.1704 | 18.8262 | 13.0908 | 16.9319 |
| Mapped | 12.0193 | 12.9093 | 11.8424 | 12.6621 | 12.0495 | 13.7355 | 10.2776 | 12.1704 | 18.8262 | 13.0908 | 16.9319 |

*Supplementary Table 2:* Sequencing statistics and quality for n = 11 San Marcos male samples at each stage of the data processing pipeline for raw reads as well as coverage throughout the alignment workflow. Coverage statistics are based on an alignment to *Hypostomus* reference genome. Missing data represented with "NA".

| | M1 | M2 | M4 | M5 | M6 | M7 | M8 | M9 | M10 | M11 | M12 |
|---|---|---|---|---|---|---|---|---|---|---|---|
| **R1** | | | | | | | | | | | |
| Raw (Lane 1) | 45998565 | 57989603 | 57818558 | 111679861 | 48846257 | 55013819 | 104812053 | 64538526 | 44939362 | 56849778 | 45457247 |
| Raw (Lane 2) | 45447440 | 57362011 | 57875957 | 110669912 | 48737863 | 54564082 | 103793190 | 63803646 | 45268387 | 55930652 | 45420521 |
| Raw (Lane 3) | 46177093 | 57466765 | 57978466 | 110704549 | 48578805 | 54617398 | 105402722 | 64314228 | 45231788 | 56584564 | 45310625 |
| Raw Total | 137623098 | 172818379 | 173672981 | 333054322 | 146162925 | 164195299 | 314007965 | 192656400 | 135439537 | 169364994 | 136188393 |
| Concatenated | 137623098 | 172818379 | 173672981 | NA | 146162925 | 164195299 | NA | 192656400 | 135439537 | 169364994 | 136188393 |
| CutAdapt | 137623098 | 172818379 | 173672981 | 333054322 | 146162925 | 164195299 | 314007965 | 192656400 | 135439537 | 169364994 | 136188393 |
| Trimmomatic | 123570922 | 155399646 | 158099558 | 301999780 | 133061301 | 147385367 | 284519156 | 172659814 | 121788673 | 150528728 | 118981648 |
| **R2** | | | | | | | | | | | |
| Raw (Lane 1) | 45998565 | 57989603 | 57818558 | 111679861 | 48846257 | 55013819 | 104812053 | 64538526 | 44939362 | 56849778 | 45457247 |
| Raw (Lane 2) | 45447440 | 57362011 | 57875957 | 110669912 | 48737863 | 54564082 | 103793190 | 63803646 | 45268387 | 55930652 | 45420521 |
| Raw (Lane 3) | 46177093 | 57466765 | 57978466 | 110704549 | 48578805 | 54617398 | 105402722 | 64314228 | 45231788 | 56584564 | 45310625 |
| Raw Total | 137623098 | 172818379 | 173672981 | 333054322 | 146162925 | 164195299 | 314007965 | 192656400 | 135439537 | 169364994 | 136188393 |
| Concatenated | 137623098 | 172818379 | 173672981 | NA | 146162925 | 164195299 | NA | 192656400 | 135439537 | 169364994 | 136188393 |
| CutAdapt | 137623098 | 172818379 | 173672981 | 333054322 | 146162925 | 164195299 | 314007965 | 192656400 | 135439537 | 169364994 | 136188393 |
| Trimmomatic | 123570922 | 155399646 | 158099558 | 301999780 | 133061301 | 147385367 | 284519156 | 172659814 | 121788673 | 150528728 | 118981648 |
| **Total # Reads and Base Pairs** | | | | | | | | | | | |
| Total reads | 134877674 | 169672372 | 171132360 | 327718067 | 143704209 | 161534171 | 308796138 | 189078857 | 133135853 | 166025539 | 133880302 |
| Total base pairs | 38767289400 | 48760802700 | 49384787700 | 94457677050 | 41514826500 | 46337930700 | 88997294100 | 54260800650 | 38238678900 | 47483140050 | 37929292500 |
| **Coverage** | | | | | | | | | | | |
| Pre-alignment | 19.3836447 | 24.38040135 | 24.69239385 | 47.228838525 | 20.75741325 | 23.16896535 | 44.49864705 | 27.130400325 | 19.11933945 | 23.741570025 | 18.96464625 |
| Post-alignment | 17.6936 | 22.253 | 22.4177 | 42.9117 | 18.791 | 21.0385 | 40.2934 | 24.6062 | 16.7168 | 21.5514 | 17.0265 |
| Rgadd | 17.6936 | 22.253 | 22.4177 | 42.9117 | 18.791 | 21.0385 | 40.2934 | 24.6062 | 16.7168 | 21.5514 | 17.0265 |
| Dupsmarked | 11.1401 | 14.5285 | 14.7101 | 28.4885 | 12.56583 | 13.1586 | 25.6738 | 15.8474 | 10.8892 | 13.118 | 10.0795 |
| Mapped | 11.1401 | 14.5285 | 14.7101 | 28.4885 | 12.56583 | 13.1586 | 25.6738 | 15.8474 | 10.8892 | 13.118 | 10.0795 |

*Supplementary Table 3:* Sequencing statistics and quality for n = 14 Comal female samples at each stage of the data processing pipeline for raw reads as well as coverage throughout the alignment workflow. Coverage statistics are based on an alignment to *Pterygoplichthys* reference genome. Missing data represented with "NA".

| | F1 | F2 | F3 | F4 | F5 | F6 | F7 | F8 | F9 | F10 | F11 | F12 | F13 | F14 |
|---|---|---|---|---|---|---|---|---|---|---|---|---|---|---|
| **R1** | | | | | | | | | | | | | | |
| Raw (Lane 1) | 86668034 | 49512405 | 46050250 | 53739187 | 65457545 | 56513183 | 56214474 | 57153959 | 60084664 | 71213286 | 53654869 | 61030640 | 1659516 | 52790996 |
| Raw (Lane 2) | 86338560 | 49608572 | 46586636 | 53939799 | 65732532 | 56436564 | 56849145 | 57419200 | 60821368 | 71730634 | 54804995 | 60967326 | 1621271 | 52964777 |
| Raw (Lane 3) | 260256603 | 148651760 | 139111513 | 161591978 | 196771530 | 169445073 | 170132070 | 172396265 | 181663045 | 214545063 | 162762513 | 183518175 | 4913660 | 158440016 |
| Raw Total | 260256603 | 148651760 | 139111513 | 161591978 | 196771530 | 169445073 | 170132070 | 172396265 | 181663045 | 214545063 | 162762513 | 183518175 | 4913660 | 158440016 |
| Concatenated | 260256603 | 148651760 | 139111513 | 161591978 | 196771530 | 169186329 | 170132070 | 172396265 | 181663045 | 214545063 | 162762513 | 183518175 | 4913660 | 158440016 |
| CutAdapt | 260256603 | 148651760 | 139111513 | 161591978 | 196771530 | 169186329 | 170132070 | 172396265 | 181663045 | 214545063 | 162762513 | 183518175 | 4913660 | 158440016 |
| Trimmomatic | 236693353 | 132357216 | 121794353 | 142949766 | 179037953 | 152834646 | 153750874 | 153776679 | 162433890 | 194338869 | 145063828 | 165759537 | 3244158 | 142134689 |
| **R2** | | | | | | | | | | | | | | |
| Raw (Lane 1) | 87250009 | 49530783 | 46474627 | 53912992 | 65581453 | 56495326 | 57068451 | 57823106 | 60757013 | 71601143 | 54302649 | 61520209 | 1632873 | 52684243 |
| Raw (Lane 2) | 86668034 | 49512405 | 46050250 | 53739187 | 65457545 | 56513183 | 56214474 | 57153959 | 60084664 | 71213286 | 53654869 | 61030640 | 1659516 | 52790996 |
| Raw (Lane 3) | 86338560 | 49608572 | 46586636 | 53939799 | 65732532 | 56436564 | 56849145 | 57419200 | 60821368 | 71730634 | 54804995 | 60967326 | 1621271 | 52964777 |
| Raw Total | 260256603 | 148651760 | 139111513 | 161591978 | 196771530 | 169445073 | 170132070 | 172396265 | 181663045 | 214545063 | 162762513 | 183518175 | 4913660 | 158440016 |
| Concatenated | 260256603 | 148651760 | 139111513 | 161591978 | 196771530 | 169186329 | 170132070 | 172396265 | 181663045 | 214545063 | 162762513 | 183518175 | 4913660 | 158440016 |
| CutAdapt | 260256603 | 148651760 | 139111513 | 161591978 | 196771530 | 169186329 | 170132070 | 172396265 | 181663045 | 214545063 | 162762513 | 183518175 | 4913660 | 158440016 |
| Trimmomatic | 236693353 | 132357216 | 121794353 | 142949766 | 179037953 | 152834646 | 153750874 | 153776679 | 162433890 | 194338869 | 145063828 | 165759537 | 3244158 | 142134689 |
| **Total # Reads and Base Pairs** | | | | | | | | | | | | | | |
| Total reads | 255952469 | 146420904 | 136574553 | 159103132 | 194014611 | 166624021 | 166977498 | 169158353 | 178387039 | 210636906 | 159525669 | 180251770 | 4829240 | 156002500 |
| Total base pairs | 73896873300 | 41816718000 | 38755335900 | 45307934700 | 5595788560 | 47918800050 | 48109255800 | 48440254800 | 51123139350 | 60746366250 | 45688424550 | 51901696050 | 1211009700 | 44720578350 |
| **Coverage** | | | | | | | | | | | | | | |
| Pre-alignment | 36.94843665 | 20.908359 | 19.37766795 | 22.653967 | 27.9289 | 23.9594 | 24.0546279 | 24.2201274 | 25.5615697 | 30.3731831 | 22.8442123 | 25.950848 | 0.60550485 | 22.3602892 |
| Post-alignment | 45.4356 | 25.8544 | 23.9325 | 27.7044 | 34.8116 | 29.6636 | 29.8765 | 29.9646 | 31.8259 | 37.7192 | 28.399 | 32.3052 | 0.420095 | 27.8452 |
| Rgadd | 45.4356 | 25.8544 | 23.9325 | 27.7044 | 34.8116 | 29.6636 | 29.8765 | NA | NA | 37.7192 | 28.399 | 32.3052 | 0.420095 | 27.8452 |
| Dupsmarked | 30.1479 | 15.5564 | 13.7697 | 15.9037 | 21.4324 | 18.1494 | 18.7674 | NA | NA | 23.8033 | 17.4201 | 20.8629 | 0.136571 | 16.4025 |
| Mapped | 27.8366 | 15.5564 | 13.7697 | 15.9037 | 21.4324 | 18.1494 | 18.7674 | NA | NA | 23.8033 | 17.4201 | 20.8629 | 0.136571 | 16.4025 |

*Supplementary Table 4:* Sequencing statistics and quality for n = 11 Comal male samples at each stage of the data processing pipeline for raw reads as well as coverage throughout the alignment workflow. Coverage statistics are based on an alignment to *Pterygoplichthys* reference genome. Missing data represented with "NA".

| | M1 | M2 | M3 | M4 | M5 | M6 | M7 | M8 | M9 | M10 | M11 |
|---|---|---|---|---|---|---|---|---|---|---|---|
| **R1** | | | | | | | | | | | |
| Raw (Lane 1) | 80400664 | 91124772 | 55300366 | 47676184 | 51318751 | 405392 | 61263559 | 50501246 | 86734914 | 49558232 | 59423985 |
| Raw (Lane 2) | 80457717 | 90558991 | 55643865 | 47562832 | 51206035 | 419393 | 61196837 | 50158405 | 86144998 | 49200962 | 58571925 |
| Raw (Lane 3) | 80497617 | 91778623 | 55573780 | 47627486 | 51249659 | 403952 | 61110197 | 50542523 | 87083793 | 49822342 | 59261057 |
| Raw Total | 241355998 | 273462386 | 166518011 | 142866502 | 153774445 | 1228737 | 183570593 | 151202174 | 259963705 | 148581536 | 177256967 |
| Concatenated | 241355998 | 273462386 | 166518011 | 142866502 | 153774445 | 1228737 | 183570593 | 151202174 | 259963705 | 148581536 | 177256967 |
| CutAdapt | 241355998 | 273462386 | 166518011 | 142866502 | 153774445 | 1228737 | 183570593 | 151202174 | 259963705 | 148581536 | 177256967 |
| Trimmomatic | 220029865 | 243910400 | 149379338 | 126889845 | 139313725 | 333109 | 166986267 | 134412036 | 232269895 | 131678541 | 157253050 |
| **R2** | | | | | | | | | | | |
| Raw (Lane 1) | 80400664 | 91124772 | 55300366 | 47676184 | 51318751 | 405392 | 61263559 | 50501246 | 86734914 | 49558232 | 59423985 |
| Raw (Lane 2) | 80457717 | 90558991 | 55643865 | 47562832 | 51206035 | 419393 | 61196837 | 50158405 | 86144998 | 49200962 | 58571925 |
| Raw (Lane 3) | 80497617 | 91778623 | 55573780 | 47627486 | 51249659 | 403952 | 61110197 | 50542523 | 87083793 | 49822342 | 59261057 |
| Raw Total | 241355998 | 273462386 | 166518011 | 142866502 | 153774445 | 1228737 | 183570593 | 151202174 | 259963705 | 148581536 | 177256967 |
| Concatenated | 241355998 | 273462386 | 166518011 | 142866502 | 153774445 | 1228737 | 183570593 | 151202174 | 259963705 | 148581536 | 177256967 |
| CutAdapt Out | 241355998 | 273462386 | 166518011 | 142866502 | 153774445 | 1228737 | 183570593 | 151202174 | 259963705 | 148581536 | 177256967 |
| Trimmomatic | 220029865 | 243910400 | 149379338 | 126889845 | 139313725 | 333109 | 166986267 | 134412036 | 232269895 | 131678541 | 157253050 |
| **Total # Reads and Base Pairs** | | | | | | | | | | | |
| Total reads | 237260386 | 268872119 | 164195611 | 140357385 | 151310027 | 1196868 | 180620970 | 148611051 | 255120714 | 145702190 | 173849105 |
| Total base pairs | 68593537650 | 76917377850 | 76917377850 | 40087084500 | 43593562800 | 229496550 | 52141085550 | 42453463050 | 73108591350 | 41607108650 | 49665323250 |
| **Coverage** | | | | | | | | | | | |
| Pre-alignment | 34.296768825 | 38.458688925 | 23.518121175 | 20.04354225 | 21.7967814 | 0.114748275 | 26.070542775 | 21.226731525 | 36.554295675 | 20.80354825 | 24.832661625 |
| Post-alignment | 42.3923 | 47.0671 | 29.2271 | 24.6898 | 27.17753 | 0.0125775 | 32.403 | 26.2098 | 45.164 | 25.7369 | 1.31785 |
| Rgadd | 42.3923 | 47.0671 | 29.2271 | 24.6898 | 27.17753 | 0.0125775 | 32.403 | 26.2098 | 45.164 | 25.7369 | 1.31785 |
| Dupsmarked | 27.0395 | 29.858 | 17.04 | 15.4519 | 16.8092 | < 0.01 | 20.7421 | 15.7588 | 28.645 | 16.5556 | 1.06583 |
| Mapped | 27.0395 | 27.3564 | 17.04 | 15.4519 | 16.8092 | < 0.01 | 20.7421 | 15.7588 | 28.1686 | 16.5556 | 1.06583 |

*Supplementary Table 5:* Sequencing statistics and quality for n = 12 San Felipe female samples at each stage of the data processing pipeline for raw reads as well as coverage throughout the alignment workflow. Coverage statistics are based on an alignment to the *Hypostomus* reference genome. Missing data represented with "NA".

| | F1 | F2 | F3 | F4 | F5 | F6 | F7 | F8 | F9 | F10 | F11 | F12 |
|---|---|---|---|---|---|---|---|---|---|---|---|---|
| **R1** | | | | | | | | | | | | |
| Raw (Lane 1) | 67022775 | 53927235 | 59506051 | 59875048 | 50183131 | 72560731 | 65329521 | 52447709 | 65560660 | 54401051 | 70269964 | 47874293 |
| Raw (Lane 2) | 66387825 | 53720811 | 58716375 | 59472846 | 50745573 | 72096492 | 64389399 | 52258200 | 65346446 | 53905111 | 69493976 | 46924954 |
| Raw (Lane 3) | 66908235 | 54303957 | 58396518 | 59792781 | 50947829 | 72776282 | 64839116 | 52093393 | 65469764 | 54154987 | 69897752 | 47883628 |
| Raw total | 200318835 | 161952003 | 176618944 | 179140675 | 151876533 | 217433505 | 194558036 | 156799302 | 196376870 | 162461149 | 209661692 | 142682875 |
| Concatenated | 200318835 | 161952003 | 176618944 | 179140675 | 151876533 | 217433505 | NA | 156799302 | 196376870 | 162461149 | 209661692 | 142682875 |
| CutAdapt Out (R1) | 200318835 | 161952003 | 176618944 | 179140675 | 151876533 | 217433505 | 6205863 | 156799302 | 196376870 | 162461149 | 209661692 | 142682875 |
| Trimmomatic Out (R1) | 182481253 | 147088162 | 157310094 | 163065394 | 137070217 | 196864404 | 5500806 | 141565510 | 176998469 | 147807765 | 190523066 | 127098258 |
| **R2** | | | | | | | | | | | | |
| Raw (Lane 1) | 67022775 | 53927235 | 59506051 | 59875048 | 50183131 | 72560731 | 65329521 | 52447709 | 65560660 | 54401051 | 70269964 | 47874293 |
| Raw (Lane 2) | 66387825 | 53720811 | 58716375 | 59472846 | 50745573 | 72096492 | 64389399 | 52258200 | 65346446 | 53905111 | 69493976 | 46924954 |
| Raw (Lane 3) | 66908235 | 54303957 | 58396518 | 59792781 | 50947829 | 72776282 | 64839116 | 52093393 | 65469764 | 54154987 | 69897752 | 47883628 |
| Raw total | 200318835 | 161952003 | 176618944 | 179140675 | 151876533 | 217433505 | 194558036 | 156799302 | 196376870 | 162461149 | 209661692 | 142682875 |
| Concatenated | 200318835 | 161952003 | 176618944 | 179140675 | 151876533 | 217433505 | 194558036 | 156799302 | 196376870 | 162461149 | 209661692 | 142682875 |
| CutAdapt | 200318835 | 161952003 | 176618944 | 179140675 | 151876533 | 217433505 | 194558036 | 156799302 | 196376870 | 162461149 | 209661692 | 142682875 |
| Trimmomatic | 182481253 | 147088162 | 157310094 | 163065394 | 137070217 | 196864404 | 5500806 | 141565510 | 176998469 | 147807765 | 190523066 | 127098258 |
| **Total # Reads and Base Pairs** | | | | | | | | | | | | |
| Total reads | 197123433 | 159359980 | 173565960 | 176277793 | 149359792 | 213615210 | 6087700 | 154178790 | 192832411 | 159600663 | 206379953 | 139704903 |
| Total base pairs | 56940702900 | 45967221300 | 49631408100 | 50901478050 | 42964501350 | 61571942100 | 1738275900 | 44361645000 | 55474632000 | 46111264200 | 59535452850 | 40020474150 |
| **Coverage** | | | | | | | | | | | | |
| Pre-alignment | 28.47035145 | 22.98361065 | 24.81570405 | 25.450739025 | 21.482250675 | 30.78597105 | 0.86913795 | 22.1808225 | 27.737316 | 23.0556321 | 29.767726425 | 20.010237075 |
| Post-alignment | 25.9692 | 20.9314 | 22.3956 | 23.1399 | 19.4778 | 1.00695 | 0.785636 | 20.1356 | 25.1563 | 20.8327 | 27.0586 | 18.1433 |
| Rgadd | 25.9692 | 20.9314 | 22.3956 | 23.1399 | 19.4778 | 1.00695 | 0.785636 | 20.1356 | 25.1563 | 20.8327 | 27.0586 | 18.1433 |
| Dupsmarked | 16.4975 | 13.1827 | 14.9703 | 14.8895 | 12.841 | 0.85556 | 0.575402 | 13.685 | 15.8028 | 13.8601 | 17.1272 | 11.6785 |
| Mapped | 16.4975 | 13.1827 | 14.9703 | 14.8895 | 12.841 | 0.85556 | 0.575402 | 13.685 | 15.8028 | 13.8601 | 17.1272 | 11.6785 |

*Supplementary Table 6:* Sequencing statistics and quality for n = 12 San Felipe male samples at each stage of the data processing pipeline for raw reads as well as coverage throughout the alignment workflow. Coverage statistics are based on an alignment to the *Hypostomus* reference genome.

| | **M1** | **M2** | **M3** | **M4** | **M5** | **M6** | **M7** | **M8** | **M9** | **M10** | **M11** | **M12** |
|---|---|---|---|---|---|---|---|---|---|---|---|---|
| **R1** | | | | | | | | | | | | |
| Raw (Lane 1) | 48005111 | 48075861 | 47334750 | 56989231 | 57886779 | 60957699 | 50340051 | 58815795 | 49826078 | 62228842 | 45018836 | 69585864 |
| Raw (Lane 2) | 47673790 | 47726159 | 47083943 | 56453061 | 57085890 | 60423899 | 49713203 | 58127996 | 49159683 | 61936925 | 44837685 | 69255325 |
| Raw (Lane 3) | 47899706 | 47965369 | 47300140 | 57729324 | 58210880 | 61302924 | 50394569 | 58576563 | 49760100 | 62379896 | 45147996 | 69583319 |
| Raw total | 143578607 | 143767389 | 141718833 | 171171616 | 173183549 | 182684522 | 150447823 | 175520354 | 148745861 | 186545663 | 135004517 | 208424508 |
| Concatenated | 143578607 | 143767389 | 141718833 | 171171616 | 173183549 | 182684522 | 150447823 | 175520354 | 148745861 | 186545663 | 135004517 | 208424508 |
| CutAdapt | 143578607 | 143767389 | 141718833 | 171171616 | 173183549 | 182684522 | 150447823 | 175520354 | 148745861 | 186545663 | 135004517 | 208424508 |
| Trimmomatic | 129256444 | 128834416 | 127863223 | 154927860 | 151316476 | 164284172 | 135558233 | 157982891 | 133069559 | 171010601 | 121988514 | 191560284 |
| **R2** | | | | | | | | | | | | |
| Raw (Lane 1) | 48005111 | 48075861 | 47334750 | 56989231 | 57886779 | 60957699 | 50340051 | 58815795 | 49826078 | 62228842 | 45018836 | 69585864 |
| Raw (Lane 2) | 47673790 | 47726159 | 47083943 | 56453061 | 57085890 | 60423899 | 49713203 | 58127996 | 49159683 | 61936925 | 44837685 | 69255325 |
| Raw (Lane 3) | 47899706 | 47965369 | 47300140 | 57729324 | 58210880 | 61302924 | 50394569 | 58576563 | 49760100 | 62379896 | 45147996 | 69583319 |
| Raw total | 143578607 | 143767389 | 141718833 | 171171616 | 173183549 | 182684522 | 150447823 | 175520354 | 148745861 | 186545663 | 135004517 | 208424508 |
| Concatenated | 143578607 | 143767389 | 141718833 | 171171616 | 173183549 | 182684522 | 150447823 | 175520354 | 148745861 | 186545663 | 135004517 | 208424508 |
| CutAdapt | 143578607 | 143767389 | 141718833 | 171171616 | 173183549 | 182684522 | 150447823 | 175520354 | 148745861 | 186545663 | 135004517 | 208424508 |
| Trimmomatic | 129256444 | 128834416 | 127863223 | 154927860 | 151316476 | 164284172 | 135558233 | 157982891 | 133069559 | 171010601 | 121988514 | 191560284 |
| **Total # Reads and Base Pairs** | | | | | | | | | | | | |
| Total reads | 132554894 | 141021005 | 139008767 | 168236913 | 170181103 | 179366920 | 147894626 | 172189810 | 145971136 | 183821054 | 132554894 | 205198911 |
| Total base pairs | 38181511200 | 49887193350 | 40030798500 | 48474715950 | 48224636850 | 51547663800 | 42517928850 | 49525905150 | 41856104250 | 53224748250 | 38181511200 | 59513879250 |
| **Coverage** | | | | | | | | | | | | |
| Pre-alignment | 19.0907556 | 24.943596675 | 20.01539925 | 24.237357975 | 24.11231843 | 25.7738319 | 21.25896443 | 24.762952558 | 20.92805213 | 26.61237413 | 19.0907556 | 29.75693963 |
| Post-alignment | 18.3047 | 18.2969 | 18.111 | 22.1114 | 21.6624 | 0.846384 | 19.4021 | 22.4755 | 19.0625 | 24.2161 | 17.3133 | 26.8891 |
| Rgadd | 18.3047 | 18.2969 | 18.111 | 22.1114 | 21.6624 | 0.846384 | 19.4021 | 22.4755 | 19.0625 | 24.2161 | 17.3133 | 26.8891 |
| Dupsmarked | 12.099 | 11.9963 | 11.6464 | 13.5107 | 13.5242 | 0.72086 | 12.0658 | 14.7351 | 11.961 | 15.6174 | 11.5379 | 18.0543 |
| Mapped | 12.099 | 11.9963 | 11.6464 | 13.5107 | 13.5242 | 0.72086 | 12.0658 | 14.7351 | 11.961 | 15.6174 | 11.5379 | 18.0543 |

*Supplementary Table 7:* Number of repeat sequences present in each repeat family in *Hypostomus* sp. reference genome as identified by RepeatModeler and RepeatMasker.

| | Number of repeats in assembly |
|---|---|
| **DNA** | 33 |
| **DNA/Academ-1** | 1 |
| **DNA/CMC-Chapaev-3** | 7 |
| **DNA/CMC-EnSpm** | 14 |
| **DNA/Crypton** | 4 |
| **DNA/Crypton-A** | 4 |
| **DNA/Crypton-V** | 1 |
| **DNA/Ginger-1** | 3 |
| **DNA/hAT** | 4 |
| **DNA/hAT-Ac** | 128 |
| **DNA/hAT-Blackjack** | 8 |
| **DNA/hAT-Charlie** | 37 |
| **DNA/hAT-hAT5** | 5 |
| **DNA/hAT-hATm** | 2 |
| **DNA/hAT-Tip100** | 19 |
| **DNA/IS3EU** | 5 |
| **DNA/Kolobok-T2** | 1 |
| **DNA/Maverick** | 3 |
| **DNA/MULE-MuDR** | 1 |
| **DNA/P** | 2 |
| **DNA/PIF** | 1 |
| **DNA/PIF-Harbinger** | 40 |
| **DNA/PIF-ISL2EU** | 4 |
| **DNA/PiggyBac** | 11 |
| **DNA/Sola-1** | 1 |
| **DNA/Sola-2** | 1 |

| | |
|---|---|
| **DNA/TcMar** | 2 |
| **DNA/TcMar-Fot1** | 3 |
| **DNA/TcMar-ISRm11** | 6 |
| **DNA/TcMar-Mariner** | 1 |
| **DNA/TcMar-Tc1** | 187 |
| **DNA/TcMar-Tc2** | 2 |
| **DNA/TcMar-Tigger** | 75 |
| **LINE/I** | 25 |
| **LINE/L1** | 20 |
| **LINE/L1-Tx1** | 8 |
| **LINE/L2** | 68 |
| **LINE/Penelope** | 13 |
| **LINE/R2-Hero** | 6 |
| **LINE/R2-NeSL** | 1 |
| **LINE/Rex-Babar** | 50 |
| **LINE/RTE-BovB** | 18 |
| **LINE/RTE-X** | 11 |
| **LTR** | 5 |
| **LTR/Copia** | 1 |
| **LTR/ERV1** | 22 |
| **LTR/ERVK** | 1 |
| **LTR/Gypsy** | 46 |
| **LTR/Ngaro** | 17 |
| **LTR/Pao** | 6 |
| **RC/Helitron** | 7 |
| **Retroposon** | 1 |
| **rRNA** | 2 |
| **Satellite** | 6 |

| | |
|---|---|
| **Simple_repeat** | 24 |
| **SINE?** | 2 |
| **SINE/5S** | 1 |
| **SINE/B2** | 2 |
| **SINE/ID** | 5 |
| **SINE/MIR** | 6 |
| **SINE/tRNA** | 4 |
| **SINE/tRNA-RTE** | 5 |
| **SINE/tRNA-V-RTE** | 1 |
| **tRNA** | 13 |
| **Unknown** | 1295 |

*Supplementary Table 8:* List of cytochrome oxidase I (COI) gene sequences from Genbank used for mitochondrial phylogeny.

| Accession No. | Taxon |
|---|---|
| AP012021.1 | *Pterygoplichthys disjunctivus* |
| EU359422.1 | *Hypostomus boulengeri* |
| FJ434521.1 | *Hypostomus ancistroides* |
| GQ225397.1 | *Hypostomus cochliodon* |
| GU701547.1 | *Pterygoplichthys ambrosettii* |
| GU701550.1 | *Pterygoplichthys ambrosettii* |
| GU701551.1 | *Pterygoplichthys ambrosettii* |
| GU701680.1 | *Hypostomus regani* |
| GU701683.1 | *Hypostomus paulinus* |
| GU701684.1 | *Hypostomus paulinus* |
| GU701687.1 | *Hypostomus paulinus* |
| GU701688.1 | *Hypostomus nigromaculatus* |
| GU701689.1 | *Hypostomus nigromaculatus* |
| GU701691.1 | *Hypostomus* sp. |
| GU701692.1 | *Hypostomus myersi* |
| GU701693.1 | *Hypostomus myersi* |
| GU701694.1 | *Hypostomus microstomus* |
| GU701695.1 | *Hypostomus regani* |
| GU701696.1 | *Hypostomus myersi* |
| GU701697.1 | *Hypostomus* sp. |
| GU701698.1 | *Hypostomus microstomus* |
| GU701699.1 | *Hypostomus iheringii* |
| GU701700.1 | *Hypostomus topavae* |
| GU701705.1 | *Hypostomus heraldoi* |
| GU701706.1 | *Hypostomus hermanni* |
| GU701707.1 | *Hypostomus hermanni* |
| GU701708.1 | *Hypostomus heraldoi* |
| GU701709.1 | *Hypostomus heraldoi* |

| Accession No. | Taxon |
| --- | --- |
| GU701710.1 | *Hypostomus heraldoi* |
| GU701711.1 | *Hypostomus heraldoi* |
| GU701713.1 | *Hypostomus derbyi* |
| GU701715.1 | *Hypostomus brevis* |
| GU701717.1 | *Hypostomus commersoni* |
| GU701719.1 | *Hypostomus commersoni* |
| GU701720.1 | *Hypostomus commersoni* |
| GU701721.1 | *Hypostomus brevis* |
| GU701722.1 | *Hypostomus commersoni* |
| GU701724.1 | *Hypostomus brevis* |
| GU701725.1 | *Hypostomus ancistroides* |
| GU701726.1 | *Hypostomus ancistroides* |
| GU701727.1 | *Hypostomus ancistroides* |
| GU701728.1 | *Hypostomus albopunctatus* |
| GU701729.1 | *Hypostomus ancistroides* |
| GU701730.1 | *Hypostomus albopunctatus* |
| GU701731.1 | *Hypostomus albopunctatus* |
| GU701732.1 | *Hypostomus regani* |
| GU701877.1 | *Hypostomus strigaticeps* |
| GU701953.1 | *Hypostomus strigaticeps* |
| GU701954.1 | *Hypostomus strigaticeps* |
| GU701963.1 | *Hypostomus regani* |
| GU701966.1 | *Hypostomus regani* |
| GU702244.1 | *Hypostomus affinis* |
| GU702253.1 | *Hypostomus affinis* |
| GU702254.1 | *Hypostomus affinis* |
| GU702261.1 | *Hypostomus auroguttatus* |
| GU702262.1 | *Hypostomus auroguttatus* |
| HM376401.1 | *Hypostomus cochliodon* |

| Accession No. | Taxon |
|---|---|
| HM405132.1 | *Hypostomus alatus* |
| HM405133.1 | *Hypostomus alatus* |
| HM405134.1 | *Hypostomus alatus* |
| HM405209.1 | *Pterygoplichthys etentaculatus* |
| HM405210.1 | *Pterygoplichthys etentaculatus* |
| HQ573337.1 | *Pterygoplichthys disjunctivus* |
| HQ573338.1 | *Pterygoplichthys sp.* |
| HQ600829.1 | *Hypostomus alatus* |
| HQ682719.1 | *Pterygoplichthys pardalis* |
| HQ682720.1 | *Pterygoplichthys pardalis* |
| JF498719.1 | *Pterygoplichthys disjunctivus* |
| JF498720.1 | *Pterygoplichthys disjunctivus* |
| JF498721.1 | *Pterygoplichthys disjunctivus* |
| JF498722.1 | *Pterygoplichthys disjunctivus* |
| JF498723.1 | *Pterygoplichthys disjunctivus* |
| JF498724.1 | *Pterygoplichthys disjunctivus* |
| JF498754.1 | *Pterygoplichthys pardalis* |
| JF769356.1 | *Pterygoplichthys disjunctivus* |
| JF769357.1 | *Pterygoplichthys pardalis* |
| JF769358.1 | *Pterygoplichthys pardalis* |
| JF769359.1 | *Pterygoplichthys pardalis* |
| JF769360.1 | *Pterygoplichthys pardalis* |
| JF769361.1 | *Pterygoplichthys pardalis* |
| JF769362.1 | *Pterygoplichthys pardalis* |
| JN026849.1 | *Hypostomus plecostomus* |
| JN026850.1 | *Hypostomus plecostomus* |
| JN026851.1 | *Hypostomus plecostomus* |
| JN028316.1 | *Pterygoplichthys* cf. *disjunctivus* |
| JN988933.1 | *Hypostomus brevis* |

| Accession No. | Taxon |
| --- | --- |
| JN988934.1 | *Hypostomus brevis* |
| JN988935.1 | *Hypostomus brevis* |
| JN988937.1 | *Hypostomus commersoni* |
| JN988938.1 | *Hypostomus commersoni* |
| JN988939.1 | *Hypostomus derbyi* |
| JN988940.1 | *Hypostomus derbyi* |
| JN988943.1 | *Hypostomus iheringii* |
| JN988944.1 | *Hypostomus iheringii* |
| JN988945.1 | *Hypostomus margaritifer* |
| JN988947.1 | *Hypostomus margaritifer* |
| JN988950.1 | *Hypostomus regani* |
| JN988952.1 | *Hypostomus topavae* |
| JQ667566.1 | *Pterygoplichthys pardalis* |
| JQ667567.1 | *Pterygoplichthys pardalis* |
| JQ667568.1 | *Pterygoplichthys pardalis* |
| JX111769.1 | *Hypostomus commersoni* |
| JX111770.1 | *Hypostomus commersoni* |
| JX111771.1 | *Hypostomus commersoni* |
| JX443490.1 | *Hypostomus ancistroides* |
| JX443491.1 | *Hypostomus ancistroides* |
| JX443494.1 | *Hypostomus strigaticeps* |
| JX443495.1 | *Hypostomus strigaticeps* |
| JX443496.1 | *Hypostomus strigaticeps* |
| JX443497.1 | *Hypostomus nigromaculatus* |
| JX443498.1 | *Hypostomus nigromaculatus* |
| JX443499.1 | *Hypostomus nigromaculatus* |
| JX443500.1 | *Hypostomus nigromaculatus* |
| JX495614.1 | *Pterygoplichthys* cf. *pardalis* |
| JX495615.1 | *Pterygoplichthys* cf. *pardalis* |

| Accession No. | Taxon |
|---|---|
| KC170030.1 | *Pterygoplichthys disjunctivus* |
| KC170031.1 | *Pterygoplichthys disjunctivus* |
| KC170032.1 | *Pterygoplichthys disjunctivus* |
| KC170033.1 | *Pterygoplichthys disjunctivus* |
| KC170034.1 | *Pterygoplichthys disjunctivus* |
| KC170035.1 | *Pterygoplichthys disjunctivus* |
| KF410695.1 | *Hypostomus plecostomus* |
| KJ720692.1 | *Hypostomus plecostomus* |
| KJ720697.1 | *Hypostomus punctatus* |
| KM104508.1 | *Hypostomus microstomus* |
| KM104509.1 | *Hypostomus ancistroides* |
| KM576100.1 | *Hypostomus plecostomus* |
| KM897223.1 | *Hypostomus regani* |
| KM897247.1 | *Hypostomus regani* |
| KM897277.1 | *Hypostomus ancistroides* |
| KM897343.1 | *Hypostomus strigaticeps* |
| KM897631.1 | *Hypostomus regani* |
| KP772577.1 | *Hypostomus carinatus* |
| KP772585.1 | *Hypostomus macushi* |
| KR491517.1 | *Pterygoplichthys multiradiatus* |
| KT239003.1 | *Pterygoplichthys anisitsi* |
| KT239004.1 | *Pterygoplichthys anisitsi* |
| KT239005.1 | *Pterygoplichthys anisitsi* |
| KT239012.1 | *Hypostomus* aff. *plecostomus* |
| KT239013.1 | *Hypostomus affinis* |
| KT239016.1 | *Pterygoplichthys pardalis* |
| KT952445.1 | *Hypostomus* sp. |
| KU288820.1 | *Pterygoplichthys anisitsi* |
| KU568996.1 | *Pterygoplichthys* sp. |

| Accession No. | Taxon |
|---|---|
| KU568997.1 | *Pterygoplichthys gibbiceps* |
| KU568997.1 | *Pterygoplichthys gibbiceps* |
| KX087171.1 | *Hypostomus* sp. |
| KX087181.1 | *Pterygoplichthys* sp. |
| KY349029.1 | *Hypostomus jaguribensis* |
| KY349030.1 | *Hypostomus jaguribensis* |
| KY349037.1 | *Hypostomus pusarum* |
| KY349038.1 | *Hypostomus pusarum* |
| KY349039.1 | *Hypostomus* sp. |
| KY349040.1 | *Hypostomus* sp. |
| KY349041.1 | *Hypostomus* sp. |
| KY349042.1 | *Hypostomus* sp. |
| MG825006.1 | *Hypostomus affinis* |
| MG825026.1 | *Hypostomus luetkeni* |
| MH087048.1 | *Pterygoplichthys anisitsi* |
| MK026008.1 | *Hypostomus francisci* |
| MK049441.1 | *Pterygoplichthys anisitsi* |
| MK355286.1 | *Hypostomus* sp. |
| MK464029.1 | *Hypostomus* sp. |
| MK464032.1 | *Hypostomus* sp. |
| MK464054.1 | *Hypostomus* sp. |
| MK464060.1 | *Hypostomus* sp. |
| MK464075.1 | *Hypostomus* sp. |
| MK464108.1 | *Hypostomus* sp. |
| MK464112.1 | *Hypostomus* sp. |
| MK464113.1 | *Hypostomus* sp. |
| MK464114.1 | *Hypostomus* sp. |
| MK464115.1 | *Hypostomus* sp. |
| MK464116.1 | *Hypostomus* sp. |

| Accession No. | Taxon |
| --- | --- |
| MK464117.1 | *Hypostomus* sp. |
| MK464118.1 | *Hypostomus* sp. |
| MK464119.1 | *Hypostomus* sp. |
| MK464120.1 | *Hypostomus* sp. |
| MK464141.1 | *Hypostomus* sp. |
| MK464142.1 | *Hypostomus* sp. |
| MK464143.1 | *Hypostomus* sp. |
| MK464144.1 | *Hypostomus* sp. |
| MK464145.1 | *Hypostomus* sp. |
| MK464146.1 | *Hypostomus* sp. |
| MK464147.1 | *Hypostomus* sp. |
| MK464163.1 | *Hypostomus* sp. |
| MK464172.1 | *Hypostomus* sp. |
| MK572527.1 | *Pterygoplichthys* sp. |
| MK628372.1 | *Pterygoplichthys anisitsi* |
| MK660623.1 | *Hypostomus variipictus* |
| MK660624.1 | *Hypostomus variipictus* |
| MK959837.1 | *Pterygoplichthys scrophus* |
| MK959838.1 | *Hypostomus microstomus* |
| MK959839.1 | *Hypostomus cochliodon* |
| MK959840.1 | *Hypostomus arecuta* |
| MK959841.1 | *Hypostomus luteomaculatus* |
| MK959843.1 | *Hypostomus regani* |
| MK959844.1 | *Hypostomus paranensis* |
| MK959845.1 | *Hypostomus formosae* |
| MK959846.1 | *Hypostomus boulengeri* |
| MK959847.1 | *Hypostomus* sp. |
| MK959848.1 | *Hypostomus borellii* |
| MK959849.1 | *Hypostomus ericae* |

| Accession No. | Taxon |
|---|---|
| MK959850.1 | *Hypostomus* sp. |
| MK959851.1 | *Hypostomus interruptus* |
| MK959852.1 | *Hypostomus* sp. |
| MK959853.1 | *Hypostomus affinis* |
| MK959854.1 | *Hypostomus asperatus* |
| MK959855.1 | *Hypostomus* sp. |
| MK959856.1 | *Hypostomus* sp. |
| MK959858.1 | *Hypostomus* sp. |
| MK959859.1 | *Hypostomus* sp. |
| MK959860.1 | *Hypostomus aspilogaster* |
| MK959861.1 | *Hypostomus* sp. |
| MK959862.1 | *Hypostomus watwata* |
| MK959863.1 | *Hypostomus taphorni* |
| MK959864.1 | *Hypostomus hemiurus* |
| MK959865.1 | *Hypostomus albopunctatus* |
| MK959866.1 | *Hypostomus isbrueckeri* |
| MK959867.1 | *Hypostomus* sp. |
| MK959868.1 | *Hypostomus fonchii* |
| MK959869.1 | *Hypostomus* sp. |
| MK959870.1 | *Hypostomus* sp. |
| MK959871.1 | *Hypostomus oculeus* |
| MK959872.1 | *Hypostomus* sp. |
| MK959873.1 | *Hypostomus latifrons* |
| MK959874.1 | *Hypostomus plecostomus* |
| MK959875.1 | *Hypostomus nigromaculatus* |
| MK959876.1 | *Hypostomus luteus* |
| MK959878.1 | *Pterygoplichthys multiradiatus* |
| MK959879.1 | *Aphanotorulus ammophilus* |
| MK959880.1 | *Pterygoplichthys zuliaensis* |

| Accession No. | Taxon |
|---|---|
| MK959881.1 | *Hypostomus laplatae* |
| MK959882.1 | *Hypostomus spiniger* |
| MK959884.1 | *Hypostomus myersi* |
| MK959885.1 | *Hypostomus derbyi* |
| MK959886.1 | *Hypostomus uruguayensis* |
| MK959887.1 | *Hypostomus ternetzi* |
| MK959888.1 | *Hypostomus hondae* |
| MK959889.1 | *Hypostomus plecostomoides* |
| MK959890.1 | *Hypostomus ancistroides* |
| MK959891.1 | *Hypostomus mutucae* |
| MK959892.1 | *Hypostomus* sp. |
| MK959893.1 | *Hypostomus ancistroides* |
| MN854452.1 | *Hypostomus* sp. |
| MN854453.1 | *Hypostomus scabriceps* |
| MN854454.1 | *Hypostomus* cf. *borellii* |
| MN854455.1 | *Hypostomus taphorni* |
| MN854456.1 | *Hypostomus boulengeri* |
| MN854457.1 | *Hypostomus hemiurus* |
| MN854458.1 | *Hypostomus* sp. |
| MN854459.1 | *Hypostomus micromaculatus* |
| MN854460.1 | *Hypostomus* sp. |
| MN854461.1 | *Hypostomus* sp. |
| MN854462.1 | *Hypostomus paranensis* |
| MN854463.1 | *Hypostomus basilisko* |
| MN854464.1 | *Hypostomus* cf. *borellii* |
| MN854465.1 | *Hypostomus* sp. |
| MN854466.1 | *Hypostomus* sp. |
| MN854467.1 | *Hypostomus taphorni* |
| MN854468.1 | *Hypostomus commersoni* |

| Accession No. | Taxon |
|---|---|
| MN854469.1 | *Hypostomus commersoni* |
| MN854470.1 | *Hypostomus* sp. |
| MN854471.1 | *Hypostomus* sp. |
| MN854472.1 | *Hypostomus* sp. |
| MN854473.1 | *Hypostomus crassicauda* |
| MN854474.1 | *Hypostomus* sp. |
| MN854475.1 | *Hypostomus* sp. |
| MN854476.1 | *Hypostomus micromaculatus* |
| MN854477.1 | *Hypostomus gymnorhynchus* |
| MN854478.1 | *Hypostomus gymnorhynchus* |
| MN854479.1 | *Hypostomus gymnorhynchus* |
| MN854480.1 | *Hypostomus gymnorhynchus* |
| MN854481.1 | *Hypostomus gymnorhynchus* |
| MN854482.1 | *Hypostomus* sp. |
| MN854483.1 | *Hypostomus corantijni* |
| MN854484.1 | *Hypostomus plecostomus* |
| MN854485.1 | *Hypostomus* sp. |
| MN854486.1 | *Hypostomus plecostomus* |
| MN854487.1 | *Hypostomus* sp. |
| MN854488.1 | *Hypostomus corantijni* |
| MN854489.1 | *Hypostomus plecostomus* |
| MN854490.1 | *Hypostomus watwata* |
| MN854491.1 | *Hypostomus niceforoi* |
| MN854492.1 | *Hypostomus oculeus* |
| MN854493.1 | *Hypostomus nematopterus* |
| MN854494.1 | *Hypostomus* sp. |
| MN854495.1 | *Hypostomus mutucae* |
| MN854496.1 | *Hypostomus uruguayensis* |
| MN854497.1 | *Hypostomus faveolus* |

| Accession No. | Taxon |
|---|---|
| MN854498.1 | *Hypostomus uruguayensis* |
| MN854499.1 | *Hypostomus faveolus* |
| MN854500.1 | *Hypostomus pusarum* |
| MN854501.1 | *Aphanotorulus unicolor* |
| MN854502.1 | *Aphanotorulus unicolor* |
| MN854503.1 | *Hypostomus aspilogaster* |
| MN854504.1 | *Aphanotorulus emarginatus* |
| MN854505.1 | *Aphanotorulus emarginatus* |
| MN854506.1 | *Aphanotorulus emarginatus* |
| MN854507.1 | *Aphanotorulus emarginatus* |
| MN854508.1 | *Hypostomus hoplonites* |
| MN854509.1 | *Hypostomus* sp. |
| MN854510.1 | *Hypostomus gymnorhynchus* |
| MN854511.1 | *Hypostomus* sp. |
| MN854512.1 | *Hypostomus basilisko* |
| MN854513.1 | *Hypostomus holostictus* |
| MN854514.1 | *Hypostomus holostictus* |
| MN854515.1 | *Hypostomus robinii* |
| MN854516.1 | *Hypostomus hemicochliodon* |
| MN854517.1 | *Hypostomus plecostomoides* |
| MN854518.1 | *Hypostomus sculpodon* |
| MN854519.1 | *Hypostomus sculpodon* |
| MN854520.1 | *Hypostomus soniae* |
| MN854521.1 | *Hypostomus paucimaculatus* |
| MN854522.1 | *Aphanotorulus* sp. |
| MN854523.1 | *Hypostomus pyrineusi* |
| MN854524.1 | *Hypostomus* sp. |
| MN854525.1 | *Hypostomus* sp. |
| MN854526.1 | *Hypostomus francisci* |

| Accession No. | Taxon |
|---|---|
| MN854527.1 | *Aphanotorulus emarginatus* |
| MN854528.1 | *Aphanotorulus emarginatus* |
| MN854529.1 | *Pseudacanthicus* cf. *leopardus* |
| MN854530.1 | *Pterygoplichthys joselimaianus* |
| MN854531.1 | *Hypostomus* aff. *regani* |
| MN854532.1 | *Hypostomus plecostomus* |
| MN854533.1 | *Hypostomus plecostomus* |
| MN854534.1 | *Hypancistrus debilittera* |
| MN854535.1 | *Baryancistrus demantoides* |
| MN854536.1 | *Hypostomus* sp. |
| MN854537.1 | *Peckoltia* sp. |
| MN854538.1 | *Panaque nigrolineatus* |
| MN854539.1 | *Hypostomus gymnorhynchus* |
| MN854540.1 | *Hypostomus gymnorhynchus* |
| MN854541.1 | *Panaqolus albivermis* |
| MN854542.1 | *Panaqolus* sp. |
| MN854543.1 | *Parancistrus nudiventris* |
| MN854544.1 | *Aphanotorulus* sp. |
| MN854545.1 | *Hypostomus topavae* |
| MN854546.1 | *Pterygoplichthys* aff. *ambrosettii* |
| MN854547.1 | *Hypostomus* sp. |
| MN854548.1 | *Hypostomus* sp. |
| MN854549.1 | *Hypostomus meleagris* |
| MN854550.1 | *Hypostomus oculeus* |
| MN854551.1 | *Pterygoplichthys* aff. *ambrosettii* |
| MN854552.1 | *Pterygoplichthys* aff. *ambrosettii* |
| MN854553.1 | *Hypostomus* sp. |
| MN854554.1 | *Aphanotorulus emarginatus* |
| MN854555.1 | *Pterygoplichthys scrophus* |

| Accession No. | Taxon |
|---|---|
| MN854556.1 | *Pterygoplichthys pardalis* |
| MN854557.1 | *Isorineloricaria spinosissima* |
| MN854558.1 | *Peckoltia braueri* |
| MN854559.1 | *Peckoltia sabaji* |
| MN854560.1 | *Hypostomus auroguttatus* |
| MN854561.1 | *Hypostomus scabriceps* |
| MN854562.1 | *Pterygoplichthys gibbiceps* |
| MN854563.1 | *Hypostomus coppenamensis* |
| MN854564.1 | *Hypostomus coppenamensis* |
| MN854565.1 | *Hypostomus watwata* |
| MN854566.1 | *Peckoltia cavatica* |
| MN854567.1 | *Peckoltia vittata* |
| MN854568.1 | *Panaque bathyphilus* |
| MN854569.1 | *Hypostomus* sp. |
| MN854570.1 | *Pterygoplichthys pardalis* |
| MN854571.1 | *Hypostomus arecuta* |
| MN854572.1 | *Pterygoplichthys lituratus* |
| MN854573.1 | *Pterygoplichthys ambrosettii* |
| MN854574.1 | *Hypostomus saramaccensis* |
| MN854575.1 | *Peckoltia oligospila* |
| MN854576.1 | *Hypostomus scabriceps* |
| MN854577.1 | *Hypostomus watwata* |
| MN854578.1 | *Scobinancistrus* cf. *pariolispos* |
| MN854579.1 | *Hypostomus* aff. *weberi* |
| MN854580.1 | *Hypostomus macushi* |
| MN854581.1 | *Hypostomus macushi* |
| MN854582.1 | *Hypostomus macushi* |
| MN854583.1 | *Hypostomus hoplonites* |
| MN854584.1 | *Hypostomus crassicauda* |

| Accession No. | Taxon |
|---|---|
| MN854585.1 | *Aphanotorulus emarginatus* |
| MN854586.1 | *Hypostomus coppenamensis* |
| MN854587.1 | *Hypostomus* sp. |
| MN854588.1 | *Hypostomus saramaccensis* |
| MN854589.1 | *Hypostomus* sp. |
| MN854590.1 | *Hypostomus* sp. |
| MN854591.1 | *Hypostomus* sp. |
| MN854592.1 | *Hypostomus strigaticeps* |
| MN854593.1 | *Hypostomus pusarum* |
| MN854594.1 | *Hypostomus* sp. |
| MN854595.1 | *Hypostomus* sp. |
| MN854596.1 | *Hypostomus carinatus* |
| MN854597.1 | *Hypostomus* sp. |
| MN854598.1 | *Hypostomus* sp. |
| MN854599.1 | *Hypostomus hoplonites* |
| MN854600.1 | *Hypostomus* sp. |
| MN854601.1 | *Hypostomus* sp. |
| MN854602.1 | *Hypostomus* sp. |
| MN854603.1 | *Hypostomus* sp. |
| MN854604.1 | *Hypostomus* sp. |
| MN854605.1 | *Hypostomus paulinus* |
| MN854606.1 | *Hypostomus* sp. |
| MN854607.1 | *Hypostomus carinatus* |
| MN854608.1 | *Hypostomus weberi* |
| MN854609.1 | *Hypostomus* aff. *pyrineusi* |
| MN854610.1 | *Hypostomus weberi* |
| MN854611.1 | *Hypostomus carinatus* |
| MN854612.1 | *Hypostomus goyazensis* |
| MN854613.1 | *Hypostomus ericae* |

| Accession No. | Taxon |
|---|---|
| MN854614.1 | *Hypostomus* sp. |
| MN854615.1 | *Hypostomus paucipunctatus* |
| MN854616.1 | *Hypostomus* sp. |
| MN854617.1 | *Hypostomus* sp. |
| MN854618.1 | *Hypostomus iheringii* |
| MN854619.1 | *Hypostomus kopeyaka* |
| MN854620.1 | *Hypostomus carinatus* |
| MN854621.1 | *Hypostomus plecostomoides* |
| MN854622.1 | *Hypostomus* sp. |
| MN854623.1 | *Hypostomus weberi* |
| MN854624.1 | *Hypostomus rhantos* |
| MN854625.1 | *Hypostomus spiniger* |
| MN854626.1 | *Hypostomus* sp. |
| MN854627.1 | *Hypostomus* sp. |
| MN854628.1 | *Ancistomus snethlageae* |
| MN854629.1 | *Hypostomus* sp. |
| MN854630.1 | *Hypostomus plecostomus* |
| MN854631.1 | *Hypostomus* aff. *piratatu* |
| MN854632.1 | *Hypostomus* sp. |
| MN854633.1 | *Hypostomus albopunctatus* |
| MN880037.1 | *Hypostomus ancistroides* |
| MN880038.1 | *Hypostomus ancistroides* |
| MN880039.1 | *Hypostomus ancistroides* |
| MN880040.1 | *Hypostomus ancistroides* |
| MN880041.1 | *Hypostomus ancistroides* |
| MN880042.1 | *Hypostomus iheringii* |
| MN880043.1 | *Hypostomus iheringii* |
| MN880044.1 | *Hypostomus albopunctatus* |
| MN880045.1 | *Hypostomus albopunctatus* |

| Accession No. | Taxon |
|---|---|
| MN880046.1 | *Hypostomus albopunctatus* |
| MN880047.1 | *Hypostomus albopunctatus* |
| MN880048.1 | *Hypostomus albopunctatus* |
| MT066232.1 | *Hypostomus ancistroides* |
| MT081402.1 | *Hypostomus* sp. |
| MT396945.1 | *Hypostomus* sp. |
| MT666008.1 | *Hypostomus watwata* |
| MT884726.1 | *Pterygoplichthys* sp. |
| MW379881.1 | *Hypostomus leucophaeus* |
| MW379882.1 | *Hypostomus jaguar* |
| MW379883.1 | *Hypostomus jaguar* |
| MW379884.1 | *Hypostomus leucophaeus* |
| MW379885.1 | *Hypostomus brevicauda* |
| MW379886.1 | *Hypostomus brevicauda* |
| MW379887.1 | *Hypostomus leucophaeus* |
| MW379888.1 | *Hypostomus leucophaeus* |
| MW379889.1 | *Hypostomus leucophaeus* |
| MW379890.1 | *Hypostomus jaguar* |
| MW379891.1 | *Hypostomus jaguar* |
| MW379892.1 | *Hypostomus wuchereri* |
| MW379893.1 | *Hypostomus* aff. *unae* |
| MW379894.1 | *Hypostomus* aff. *unae* |
| MW379895.1 | *Hypostomus* aff. *unae* |
| MW379896.1 | *Hypostomus* sp. |
| MW379897.1 | *Hypostomus* sp. |
| MW379898.1 | *Hypostomus* aff. *unae* |
| MW379899.1 | *Hypostomus* aff. *unae* |
| MW379900.1 | *Hypostomus* aff. *unae* |
| MW379901.1 | *Hypostomus* sp. |

| Accession No. | Taxon |
|---|---|
| MW379902.1 | *Hypostomus* sp. |
| MW379903.1 | *Pterygoplichthys chrysostiktos* |
| MW379904.1 | *Pterygoplichthys chrysostiktos* |
| MW379905.1 | *Hypostomus jaguar* |
| MW379906.1 | *Hypostomus jaguar* |
| MW379907.1 | *Hypostomus jaguar* |
| MW379908.1 | *Hypostomus jaguar* |
| MW379909.1 | *Pterygoplichthys chrysostiktos* |
| MW379910.1 | *Hypostomus wuchereri* |
| MW379911.1 | *Hypostomus brevicauda* |
| MW379912.1 | *Hypostomus wuchereri* |
| MW379913.1 | *Hypostomus wuchereri* |
| MW379914.1 | *Hypostomus wuchereri* |
| MW379915.1 | *Hypostomus wuchereri* |
| MW379916.1 | *Hypostomus brevicauda* |
| MW379917.1 | *Hypostomus brevicauda* |
| MW379918.1 | *Hypostomus leucophaeus* |
| MW379919.1 | *Hypostomus wuchereri* |
| MW379920.1 | *Hypostomus wuchereri* |
| MW379921.1 | *Hypostomus wuchereri* |
| MW379922.1 | *Hypostomus leucophaeus* |
| MW379923.1 | *Hypostomus wuchereri* |
| MW379924.1 | *Hypostomus wuchereri* |
| MW379925.1 | *Hypostomus wuchereri* |
| MW379926.1 | *Hypostomus brevicauda* |
| MW379927.1 | *Hypostomus brevicauda* |
| MW379928.1 | *Hypostomus wuchereri* |
| MW379929.1 | *Hypostomus wuchereri* |
| MW379930.1 | *Hypostomus wuchereri* |

| Accession No. | Taxon |
|---|---|
| MW379931.1 | *Hypostomus* sp. |
| MW379932.1 | *Hypostomus* sp. |
| MW379933.1 | *Hypostomus nigrolineatus* |
| MW379934.1 | *Hypostomus nigrolineatus* |
| MW379935.1 | *Hypostomus nigrolineatus* |
| MW379936.1 | *Hypostomus nigrolineatus* |
| MW379937.1 | *Hypostomus wuchereri* |
| MW379938.1 | *Hypostomus wuchereri* |
| MW379939.1 | *Hypostomus* aff. *unae* |
| MW379940.1 | *Hypostomus* aff. *unae* |
| MW379941.1 | *Hypostomus affinis* |
| MW379942.1 | *Pterygoplichthys* aff. *ambrosettii* |
| MW379943.1 | *Hypostomus wuchereri* |
| MW379944.1 | *Hypostomus wuchereri* |
| MW379945.1 | *Hypostomus wuchereri* |
| MW379946.1 | *Hypostomus wuchereri* |
| MW379947.1 | *Hypostomus wuchereri* |
| MW379948.1 | *Hypostomus wuchereri* |
| MW379949.1 | *Hypostomus wuchereri* |
| MW379950.1 | *Hypostomus francisci* |
| MW379951.1 | *Hypostomus francisci* |
| MW379952.1 | *Hypostomus francisci* |
| MW379953.1 | *Hypostomus francisci* |
| MW379954.1 | *Hypostomus francisci* |
| MW379955.1 | *Hypostomus francisci* |
| MW379956.1 | *Hypostomus francisci* |
| MW379957.1 | *Hypostomus* cf. *macrops* |
| MW379958.1 | *Hypostomus francisci* |
| MW379959.1 | *Hypostomus francisci* |

| Accession No. | Taxon |
| --- | --- |
| MW379960.1 | *Hypostomus francisci* |
| MW379961.1 | *Hypostomus francisci* |
| MW379962.1 | *Hypostomus velhochico* |
| MW379963.1 | *Hypostomus velhochico* |
| MW379964.1 | *Hypostomus francisci* |
| MW379965.1 | *Hypostomus francisci* |
| MW379966.1 | *Hypostomus francisci* |
| MW379967.1 | *Hypostomus francisci* |
| MW379968.1 | *Hypostomus velhochico* |
| MW379969.1 | *Hypostomus* cf. *macrops* |
| MW379970.1 | *Hypostomus* cf. *macrops* |
| MW379971.1 | *Hypostomus* cf. *paulinus* |
| MW379972.1 | *Hypostomus margaritifer* |
| MW379973.1 | *Hypostomus* cf. *macrops* |
| MW379974.1 | *Hypostomus* cf. *macrops* |
| MW379975.1 | *Hypostomus* sp. |
| MW379976.1 | *Hypostomus* sp. |
| MW379977.1 | *Hypostomus* sp. |
| MW379978.1 | *Hypostomus* sp. |
| MW379979.1 | *Hypostomus* sp. |
| MW379980.1 | *Hypostomus* sp. |
| MW379981.1 | *Hypostomus* sp. |
| MW379982.1 | *Hypostomus* aff. *affinis* |
| MW379983.1 | *Hypostomus* aff. *affinis* |
| MW379984.1 | *Hypostomus* aff. *affinis* |
| MW379985.1 | *Hypostomus francisci* |
| MW379986.1 | *Hypostomus francisci* |
| MW379987.1 | *Hypostomus luetkeni* |
| MW379988.1 | *Hypostomus affinis* |

| Accession No. | Taxon |
| --- | --- |
| MW379989.1 | *Hypostomus scabriceps* |
| MW379990.1 | *Hypostomus luetkeni* |
| MW379991.1 | *Hypostomus scabriceps* |
| MW379992.1 | *Hypostomus scabriceps* |
| MW379993.1 | *Hypostomus scabriceps* |
| MW379994.1 | *Hypostomus affinis* |
| MW379995.1 | *Hypostomus affinis* |
| MW379996.1 | *Hypostomus affinis* |
| MW379997.1 | *Hypostomus affinis* |
| MW379998.1 | *Hypostomus* sp. |
| MW379999.1 | *Hypostomus francisci* |
| MW380000.1 | *Hypostomus alatus* |
| MW380001.1 | *Hypostomus luetkeni* |
| MW380002.1 | *Hypostomus affinis* |
| MW380003.1 | *Hypostomus francisci* |
| MW380004.1 | *Hypostomus* sp. |
| MW380005.1 | *Hypostomus nigrolineatus* |
| MW380006.1 | *Hypostomus* sp. |
| MW380007.1 | *Hypostomus francisci* |
| MW380008.1 | *Hypostomus alatus* |
| MW380009.1 | *Hypostomus francisci* |
| MW380010.1 | *Hypostomus* aff. *auroguttatus* |
| MW380011.1 | *Hypostomus pusarum* |
| MW380012.1 | *Hypostomus unae* |
| MW380013.1 | *Hypostomus unae* |
| MW380014.1 | *Hypostomus auroguttatus* |
| MW380015.1 | *Hypostomus affinis* |
| MW380016.1 | *Hypostomus affinis* |
| MW380017.1 | *Hypostomus jaguar* |

| Accession No. | Taxon |
|---|---|
| MW380018.1 | *Hypostomus wuchereri* |
| MW380019.1 | *Hypostomus francisci* |
| MW380020.1 | *Hypostomus francisci* |
| MW380021.1 | *Hypostomus* cf. *macrops* |
| MW380022.1 | *Hypostomus francisci* |
| MW380023.1 | *Hypostomus* aff. *francisci* |
| MW380024.1 | *Hypostomus alatus* |
| MW380025.1 | *Hypostomus alatus* |
| MW380026.1 | *Hypostomus* cf. *macrops* |
| MW380027.1 | *Hypostomus velhochico* |
| MW380028.1 | *Hypostomus velhochico* |
| MW380029.1 | *Hypostomus velhochico* |
| MW380030.1 | *Hypostomus* cf. *garmani* |
| MW380031.1 | *Hypostomus* cf. *garmani* |
| MW380032.1 | *Hypostomus* cf. *garmani* |
| MW380033.1 | *Hypostomus* cf. *garmani* |
| MW380034.1 | *Hypostomus* cf. *garmani* |
| MW380035.1 | *Hypostomus* cf. *garmani* |
| MW380036.1 | *Hypostomus* sp. |
| MW380037.1 | *Hypostomus* sp. |
| MW380038.1 | *Hypostomus* aff. *francisci* |
| MW380039.1 | *Hypostomus* aff. *francisci* |
| MW380040.1 | *Hypostomus margaritifer* |
| MW380041.1 | *Hypostomus margaritifer* |
| MW380042.1 | *Hypostomus francisci* |
| MW380043.1 | *Hypostomus* sp. |
| MW380044.1 | *Hypostomus* sp. |
| MW380045.1 | *Hypostomus leucophaeus* |
| MW380046.1 | *Hypostomus leucophaeus* |

| Accession No. | Taxon |
|---|---|
| MW380047.1 | *Hypostomus leucophaeus* |
| MW380048.1 | *Hypostomus leucophaeus* |
| MW380049.1 | *Pterygoplichthys chrysostiktos* |
| MW380050.1 | *Pterygoplichthys chrysostiktos* |
| MW380051.1 | *Hypostomus velhochico* |
| MW380052.1 | *Hypostomus velhochico* |
| MW380053.1 | *Hypostomus velhochico* |
| MW380054.1 | *Hypostomus velhochico* |
| MW380055.1 | *Hypostomus margaritifer* |
| MW380056.1 | *Hypostomus margaritifer* |
| MW380057.1 | *Hypostomus* cf. *macrops* |
| MW380058.1 | *Hypostomus* cf. *macrops* |
| MW380059.1 | *Hypostomus* aff. *francisci* |
| MW380060.1 | *Hypostomus* aff. *francisci* |
| MW380061.1 | *Hypostomus* cf. *wuchereri* |
| MW380062.1 | *Hypostomus* aff. *francisci* |
| MW401674.1 | *Pterygoplichthys pardalis* |
| MZ050829.1 | *Hypostomus plecostomus* |
| MZ050926.1 | *Hypostomus gymnorhynchus* |
| MZ050957.1 | *Hypostomus plecostomus* |
| MZ051003.1 | *Hypostomus watwata* |
| MZ051068.1 | *Hypostomus watwata* |
| MZ051282.1 | *Hypostomus gymnorhynchus* |
| MZ051559.1 | *Hypostomus plecostomus* |
| MZ051637.1 | *Hypostomus plecostomus* |
| MZ051739.1 | *Hypostomus gymnorhynchus* |
| MZ051754.1 | *Hypostomus plecostomus* |
| MZ051871.1 | *Hypostomus gymnorhynchus* |
| MZ051917.1 | *Hypostomus gymnorhynchus* |

| Accession No. | Taxon |
|---|---|
| MZ051945.1 | *Hypostomus plecostomus* |
| MZ052001.1 | *Hypostomus gymnorhynchus* |
| MZ345687.1 | *Hypostomus annectens* |
| MZ345689.1 | *Hypostomus* sp. |
| MZ345690.1 | *Hypostomus* sp. |
| MZ345691.1 | *Hypostomus* sp. |
| MZ351897.1 | *Hypostomus pusarum* |
| MZ351898.1 | *Hypostomus pusarum* |
| MZ351899.1 | *Hypostomus pusarum* |
| MZ351900.1 | *Hypostomus pusarum* |
| MZ351901.1 | *Hypostomus sertanejo* |
| MZ351902.1 | *Hypostomus sertanejo* |
| MZ351903.1 | *Hypostomus johnii* |
| MZ351904.1 | *Hypostomus johnii* |
| MZ351905.1 | *Hypostomus* sp. |
| MZ351906.1 | *Hypostomus* sp. |
| MZ351907.1 | *Hypostomus* sp. |
| MZ351908.1 | *Hypostomus* sp. |
| MZ351909.1 | *Hypostomus* sp. |
| MZ351910.1 | *Hypostomus* sp. |
| MZ351911.1 | *Hypostomus vaillanti* |
| MZ351912.1 | *Hypostomus vaillanti* |
| MZ351913.1 | *Hypostomus vaillanti* |
| MZ351914.1 | *Hypostomus vaillanti* |
| MZ351915.1 | *Hypostomus* cf. *pusarum* |
| MZ351916.1 | *Hypostomus* cf. *pusarum* |
| MZ351917.1 | *Hypostomus* cf. *pusarum* |
| MZ351918.1 | *Hypostomus* cf. *pusarum* |
| MZ351919.1 | *Hypostomus* cf. *pusarum* |

| Accession No. | Taxon |
|---|---|
| MZ351920.1 | *Hypostomus velhochico* |
| MZ351921.1 | *Hypostomus velhochico* |
| MZ351922.1 | *Hypostomus velhochico* |
| MZ351923.1 | *Parotocinclus spilurus* |
| NC_049098.1 | *Corydoras trilineatus* |
| NC_063780.1 | *Corydoras aeneus* |
| NC_063781.1 | *Corydoras paleatus* |
| NC_072072.1 | *Corydoras hastatus* |
| NC_077553.1 | *Corydoras julii* |
| NC_015747.1 | *Pterygoplichthys disjunctivus* |
| OQ200385.1 | *Hypostomus sertanejo* |
| OQ200386.1 | *Hypostomus sertanejo* |
| OQ200387.1 | *Hypostomus pusarum* |
| OQ200388.1 | *Hypostomus pusarum* |
| OQ200389.1 | *Hypostomus pusarum* |
| OQ200390.1 | *Hypostomus pusarum* |
| OQ200391.1 | *Hypostomus pusarum* |
| OQ200392.1 | *Hypostomus pusarum* |
| OQ200393.1 | *Hypostomus pusarum* |
| OQ200394.1 | *Hypostomus pusarum* |
| OQ200395.1 | *Hypostomus pusarum* |
| OQ200396.1 | *Hypostomus pusarum* |
| OQ200397.1 | *Hypostomus pusarum* |
| OQ200398.1 | *Hypostomus pusarum* |
| OQ200399.1 | *Hypostomus pusarum* |
| OQ200400.1 | *Hypostomus pusarum* |
| OQ200401.1 | *Hypostomus pusarum* |
| OQ200402.1 | *Hypostomus pusarum* |
| OQ200403.1 | *Hypostomus pusarum* |

| Accession No. | Taxon |
|---|---|
| OQ200404.1 | *Hypostomus pusarum* |
| OQ200405.1 | *Hypostomus pusarum* |
| OQ200406.1 | *Hypostomus pusarum* |
| OQ200407.1 | *Hypostomus* sp. |
| OQ200408.1 | *Hypostomus* sp. |
| OQ200409.1 | *Hypostomus* sp. |
| OQ200410.1 | *Hypostomus* sp. |
| OQ200411.1 | *Hypostomus* sp. |
| OQ200412.1 | *Hypostomus* sp. |
| OQ200413.1 | *Hypostomus* sp. |
| OQ200414.1 | *Hypostomus* sp. |
| OQ200415.1 | *Hypostomus* sp. |
| OQ200416.1 | *Hypostomus* sp. |
| OQ200417.1 | *Hypostomus* sp. |
| OQ200418.1 | *Hypostomus pusarum* |
| OQ231490.1 | *Hypostomus plecostomus* |
| OQ231491.1 | *Hypostomus plecostomus* |
| OQ231492.1 | *Hypostomus plecostomus* |
| OR732968.1 | *Hypostomus cochliodon* |
| PP998994.1 | *Hypostomus khimaera* |
| PP998995.1 | *Hypostomus khimaera* |
| PP998996.1 | *Hypostomus khimaera* |
| PP998997.1 | *Hypostomus khimaera* |
| PP998998.1 | *Hypostomus khimaera* |
| PP998999.1 | *Hypostomus khimaera* |
| PP999000.1 | *Hypostomus khimaera* |
| PQ233434.1 | *Hypostomus* sp. |
| PQ233441.1 | *Hypostomus hondae* |
| PQ233442.1 | *Hypostomus hondae* |

| Accession No. | Taxon |
|---|---|
| PQ233580.1 | *Hypostomus* sp. |
| PQ233585.1 | *Hypostomus* sp. |
| PQ233596.1 | *Hypostomus hondae* |
| PQ233603.1 | *Hypostomus* sp. |
| PQ233605.1 | *Hypostomus* sp. |
| PQ233607.1 | *Hypostomus* sp. |
| PQ233660.1 | *Hypostomus hondae* |
| PQ233702.1 | *Hypostomus hondae* |
| PQ233846.1 | *Hypostomus hondae* |
| PV077306.1 | *Hypostomus ancistroides* |
| PV077307.1 | *Hypostomus ancistroides* |
| PV077308.1 | *Hypostomus ancistroides* |
| PV077309.1 | *Hypostomus ancistroides* |
| PV077310.1 | *Hypostomus ancistroides* |
| PV077311.1 | *Hypostomus ancistroides* |
| PV077312.1 | *Hypostomus ancistroides* |
| PV077313.1 | *Hypostomus ancistroides* |
| PV077314.1 | *Hypostomus ancistroides* |
| PV077315.1 | *Hypostomus ancistroides* |
| PV077316.1 | *Hypostomus ancistroides* |
| PV077317.1 | *Hypostomus ancistroides* |
| PV077318.1 | *Hypostomus ancistroides* |
| PV241661.1 | *Hypostomus plecostomus* |
| PV241662.1 | *Hypostomus plecostomus* |

*Supplementary Table 9:* Cross-validation error rate for K = 1 through K = 3 for all-population and "true" *Hypostomus*-only ADMIXTURE analyses. The lowest value indicates the optimal number of clusters representing genetic variation in the samples.

| ADMIXTURE analysis | Cross-validation (CV) error |
|---|---|
| **All-population** | |
| K=1 | 0.94301 |
| K=2 | 0.46520 |
| K=3 | 0.42119 |
| K=4 | 0.44448 |
| K=5 | 0.58914 |
| **San Marcos River and San Felipe Creek populations only** | |
| K=1 | 0.62623 |
| K=2 | 0.56252 |
| K=3 | 0.64117 |
| K=4 | 0.70639 |
| K=5 | 0.78484 |

*Supplementary Figure 1:* IQ-TREE2 analysis showing a phylogenetic hypothesis of the mitochondrial gene cytochrome c oxidase I (COI). Samples from the San Marcos River, San Felipe Creek, and Comal River in our study are highlighted in green.

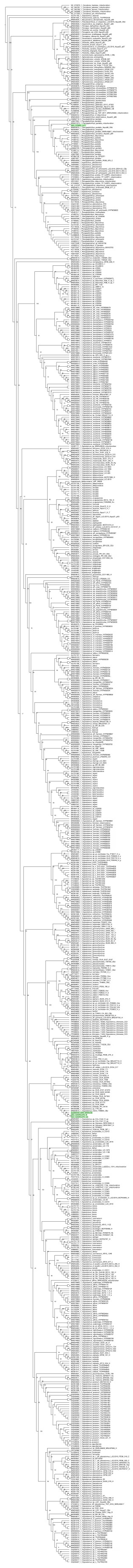

*Supplementary Figure 2:* ADMIXTURE results (K = 1–5) showing genetic population structure of suckermouth armored catfish in (a) all three Texas rivers and (b) only "true" Hypostomus species, indicating evidence of historical gene flow.

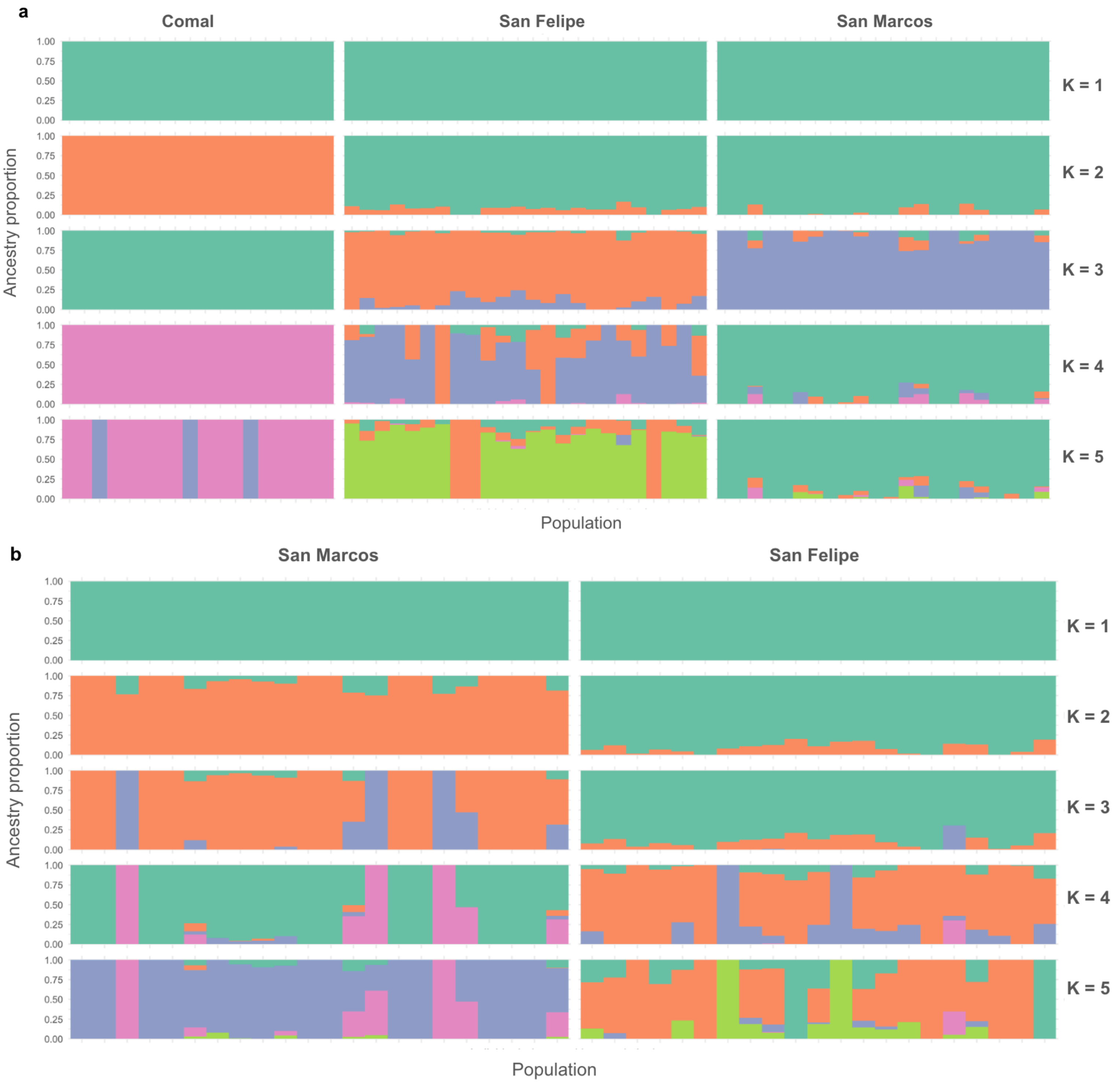
